# Exploration-based learning of a stabilizing controller predicts locomotor adaptation

**DOI:** 10.1101/2021.03.18.435986

**Authors:** Nidhi Seethapathi, Barrett Clark, Manoj Srinivasan

## Abstract

Humans adapt their locomotion seamlessly in response to changes in the body or the environment. We do not understand how such adaptation improves performance measures like energy consumption or symmetry while avoiding falling. Here, we model locomotor adaptation as interactions between a stabilizing controller that reacts quickly to perturbations and a reinforcement learner that gradually improves the controller’s performance through local exploration and memory. This model predicts time-varying adaptation in many settings: walking on a split-belt treadmill (i.e. with both feet at different speeds), with asymmetric leg weights, or using exoskeletons — capturing learning and generalization phenomena in ten prior experiments and two model-guided experiments conducted here. The performance measure of energy minimization with a minor cost for asymmetry captures a broad range of phenomena and can act alongside other mechanisms such as reducing sensory prediction error. Such a model-based understanding of adaptation can guide rehabilitation and wearable robot control.

## Introduction

Humans readily adapt their locomotion to diverse environmental conditions and bodily changes^1–3^ (Fig. 1a), but the computational principles underlying such adaptation are not fully understood. While crucial adaptation phenomena have been uncovered through careful experiments^2,3,3–8^ and a handful of models have been proposed to explain individual experiments^2,9,10^, an integrative understanding of adaptation across paradigms and timescales is missing. Moreover, existing adaptation models are not implemented on a bipedal physics-based agent, and therefore do not encompass the stability-critical nature of adapting locomotion while avoiding falling. In this work, we put forth an integrative model of locomotor adaptation combining stabilizing control, performance-improving reinforcement learning, and performance-based memory updates. Our model predicts locomotor adaptation phenomena across paradigms in ten prior studies and two prospective experiments conducted in this study.

**Figure 1.**
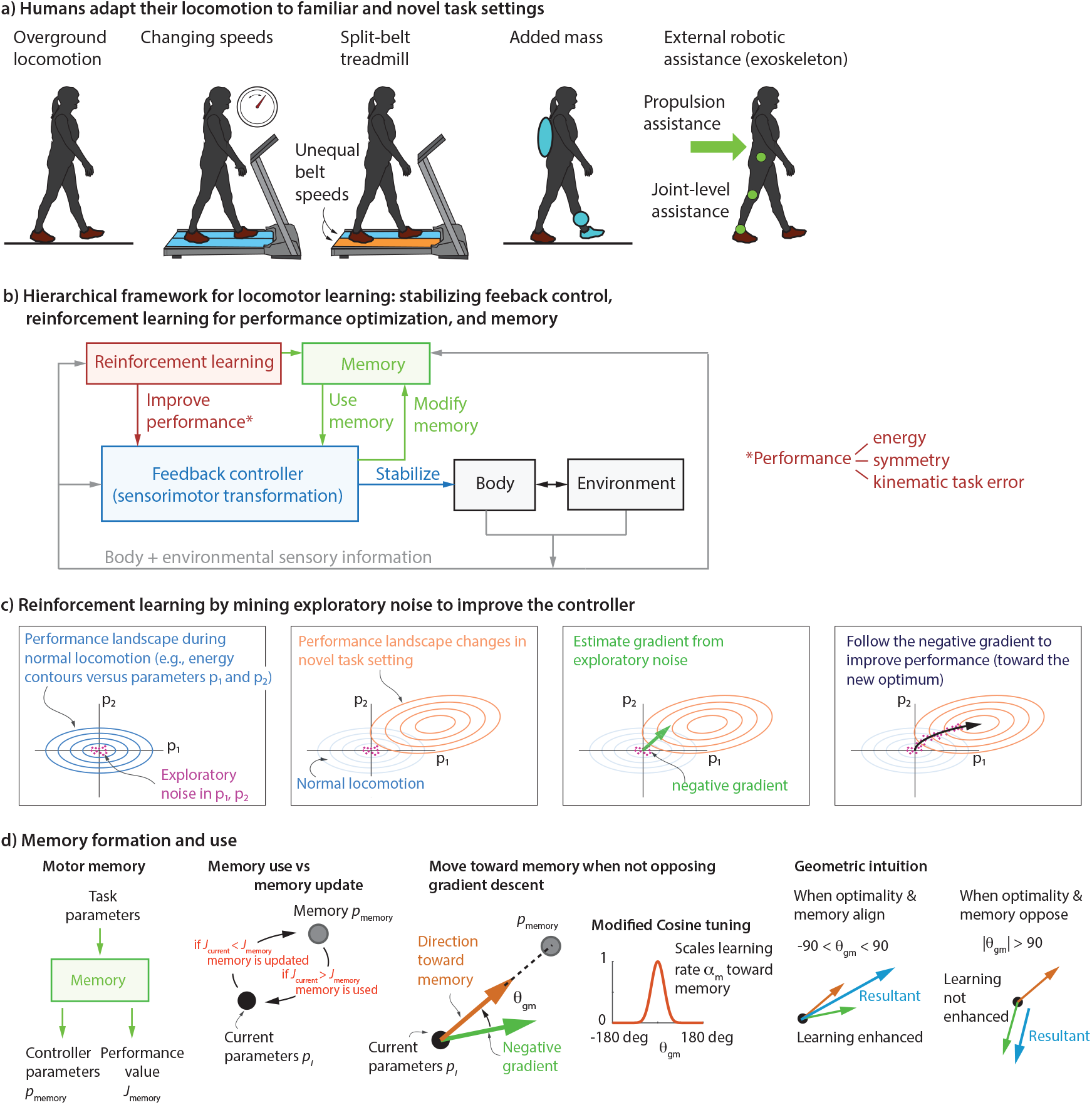
A hierarchical framework for locomotor adaptation. **a**. Humans are able to adapt readily to numerous locomotor task settings, both familiar and novel. **b**. Description of the proposed hierarchical framework, containing three components: (i) the inner loop, represents a fast timescale response due to the stabilizing feedback controller (blue), aimed at avoiding falling; (ii) an outer loop, represents reinforcement learning (red) that tunes the parameters of the inner loop controller to improve some performance objective; (iii) storing and using memories of the learned controllers (green). Alternative adaptation mechanisms may include different performance objectives within the same framework (energy, symmetry, task error) or may replace the feedback controller by a sensorimotor transformation with a state estimator followed by the controller (Fig. 9). **c**. Reinforcement learning by mining exploratory noise to estimate gradient and improve the controller. Initially, the controller parameters *p*_1_ and *p*_2_ are near the optimum of the initial performance landscape (blue). When conditions change, the performance contours change (blue to orange) as does the optimum. Exploratory noise in the controller parameters, allows the learner to estimate the gradient of the performance objective and follow the negative of this gradient to improve performance. **d**. Memory takes in task parameters and returns the stored controller parameters *p*_memory_ and the associated performance value *J*_memory_. We describe how memory is used in concert with gradient-based learning. The control parameters *p*_*i*_ are updated toward memory *p*_memory_ when doing so improves performance (memory use); memory is updated toward the current parameters otherwise. Updates toward memory is degraded if these updates are not aligned with the gradient, and this degradation is mediated by a modified cosine tuning.

Theories of motor adaptation have predominantly been developed for discrete episodic tasks such as reaching with the arm^11–13^. Adaptation principles that explain such episodic tasks may not be sufficient for explaining continuously cascading stability-critical tasks such as locomotion, multi-fingered manipulation, and many activities of daily living. In episodic tasks like reaching where the arm’s state is re-set at the end of each episode, the errors during one episode do not dynamically propagate to the next episode. In contrast, in continuously cascading tasks like locomotion, errors can have short-term and long-term consequences to stability unless otherwise controlled^14–17^. Prior accounts of locomotor adaptation^2,9,18^ do not consider the interaction with locomotor dynamics, perhaps assuming that dynamic stability is ensured by a distinct mechanism. For instance, metabolic energy reduction-based accounts^2,19^ treated adaptation to be a univariate optimization process – implicitly assuming that changes on one step do not affect the next step through the dynamics. Similarly, error-based learning models developed for arm reaching^11–13^, when applied to locomotor adaptation^9,10,18,20,21^, do not usually interact with the locomotor dynamics; these models fit the kinematic symmetry error transients, without considering how these errors might affect stability. Here, we put forth a model that explains how humans adapt continuously during walking while maintaining dynamic stability.

Improving some aspect of performance is a driving force for motor adaptation and learning. However, we do not understand which performance objectives explain diverse locomotor adaptation phenomena. Minimization of different types of error^11,12,22–24^ (e.g., sensory prediction error, task error, proprioceptive conflict) or minimization of metabolic energy^2,7,8,25^ have been separately posited as performance objectives underlying locomotor adaptation. However, these performance objectives often do not generalize across settings. Metabolic energy minimization can explain steady state adaptation in some experimental settings^25,26^ but does not in other settings^27^. Similarly, while error-based learning models can be fit to asymmetry changes in some tasks^3,4,6,10,18,24^, they cannot make predictions for tasks where there is no changes in the symmetry^28,29^. A computational model that precisely specifies the performance objectives such as energy, sensory prediction error, proprioceptive conflict, etc. would help identify the performance objectives that predict locomotor adaptation phenomena across tasks. Here, we put forth such a predictive model of locomotor adaptation allowing comparisons between the predictive ability of difference performance objectives, finding that energy minimization predicts the broadest range of phenomena.

In this work, we contribute a model of adaptation which causally links the body dynamics, stabilizing control policy, learning algorithm, performance goal, internal model of performance, and memory of control. We model adaptation as an exploration-driven gradient-based improvement of a stabilizing controller, explaining how humans improve their locomotor performance continuously while maintaining stability. Our model predicts adaptation phenomena in ten prior experimental studies and two model-guided experiments conducted here. The model captures learning phenomena such as fast timescale response followed by slow timescale adaptation, savings, faster de-adaptation, generalization, non-learning in some situations, and the effect of noise and prior experience. By modifying the performance objective, we show that our modeling framework can help compare theories of locomotor adaptation such as minimizing energy, sensory prediction error (via proprioceptive realignment), or kinematic task error (e.g., asymmetry) in their ability to explain phenomena.

## Results

### A modular and hierarchical model of locomotor adaptation

We posit a modular and hierarchical model of locomotor adaptation (Fig. 1b-d) in which a controller keeps the human stable, a gradient-based reinforcement learner modifies this stabilizing controller to improve performance, an internal model learns to predict performance in a new environment, and a memory mechanism stores the improved walking strategies and deploys them when advantageous. The model is modular in that there are separate but interacting modules performing distinct tasks (stabilizing control, gradient estimation, gradient-based learning, memory update); the model is hierarchical in that some modules operate at and explain phenomena at distinct timescales that are hierarchically separated. We test the ability of the computational model to predict experimentally observed locomotor adaptation phenomena in a number of experiments: see our repository *LocAd*^30^ for the code implementing the model.

A critical constraint on human locomotion is being stable i.e. not falling down, despite internal and external perturbations. Thus, a stabilizing controller forms the inner-most level of our hierarchical model^16,17,31^ (Fig. 1b). We posit that during locomotion in a familiar setting, humans use a previously learned controller, which we call a ‘default controller,’ stored as a motor memory. We further posit that the structure of this default controller constrains how humans adapt to a novel situation. We characterized this default controller by modeling how humans respond to small deviations from nominal walking on the treadmill^16,17,31^. This controller can be decomposed into a feedforward component, not dependent on the biped’s state, and other state-dependent feedback terms (see *Methods*). We used effectively the same initial default controller for all the locomotor adaptation tasks considered here (see *Methods* and *Supplementary Methods*). This is possible because the controller is robust to substantial noise and uncertainty as we have previously shown^16,17^, allowing the human to move stably in novel environments. It has been hypothesized that the nervous system chooses movements that optimize some performance objective, for instance, reducing energy expenditure^8,32–36^ or reducing left-right asymmetry^3,18,24,37^ (Fig. 1b). We posit that when faced with a novel circumstance, humans gradually change their default stabilizing controller to optimize performance. This performance improvement is achieved through gradient-based reinforcement learning in an outer loop around the stabilizing controller (Fig. 1b,c). We found that allowing the reinforcement learner to adapt just the feedforward terms of the controller, leaving the stabilizing feedback terms unchanged, is sufficient to explain the observed phenomena. The learner estimates the gradient descent direction using ‘intentional’ exploratory noise^2,13,38^ in the neighborhood of the default controller, contributing to increasing the step-to-step variability^16,17,31^. While the term ‘reinforcement learning’ has a multitude of algorithmic specifications^39^, here we use this term as shorthand for the proposed local exploration-based learning algorithm.

Motor adaptation involves memorization and retrieval of control policies. Here, we posit a module in the outer loop that forms longer-term motor memories^40,41^ of the controllers being learned, parameterized by the settings in which they were learned. This stored memory is used when encountering a setting similar to one previously encountered (Fig. 1b,d), interpolating and generalizing between settings via function approximation^39^. Stored memory is only used when it may improve performance and does not conflict with gradient descent (Fig. 1d). Conversely, stored memory is updated when the current controller’s performance is better than that of the motor memory. See *Methods* and the model’s implementation in code, *LocAd*^30^, for further details.

We have posited that the gradual modification of a stabilizing controller for performance optimization is a primary mechanism for locomotor adaptation. Adaptation may also result from other mechanisms such as recalibration to reduce sensory prediction error^11,22,42,43^. Here, we extend the aforementioned framework, showing that the model can incorporate sensory error-based adaptation mechanisms, replacing the feedback controller of Fig. 1b by a more general sensorimotor transformation (see Fig. 9 and *Methods*).

### Predicting fast and slow timescale learning in many locomotor settings

The model predicted locomotor adaptation phenomena in many different conditions, including a split-belt treadmill, an asymmetrically added leg mass, external assistance, exoskeleton-based perturbations, and abrupt treadmill speed changes (Fig. 2). For the reinforcement learner, we tested minimizing four performance objectives: only energy expenditure, only asymmetry (specifically, step length asymmetry, defined below), a weighted sum of energy and asymmetry, and a kinematic task error. For the results below, we use energy expenditure alone or energy expenditure with a small step length asymmetry penalty as the performance objective as these give qualitatively similar results, we use the latter when the performance objective is not explicitly mentioned. Minimization of other objectives are discussed in their own separate sections later.

**Figure 2.**
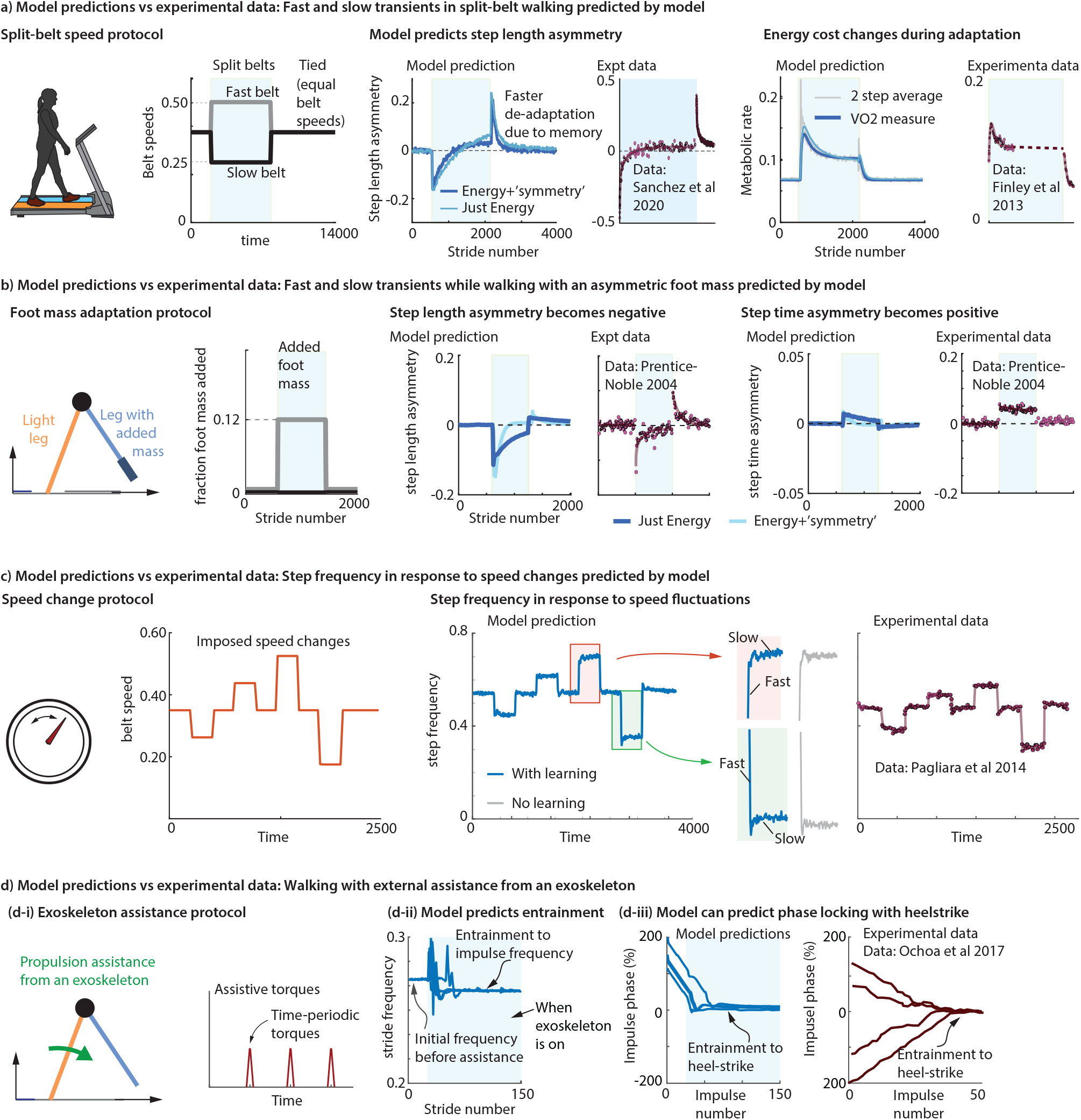
Hierarchical model predicts locomotor adaptation in multiple task settings. **a**. Split-belt walking^6,45^, that is, with the two belts going at different speeds. Model qualitatively predicts experimental transients in step length asymmetry and metabolic energy during adaptation and de-adaptation. **b**. Walking with an additional mass on one foot^5^. Model qualitatively predicts experimental transients in step length asymmetry during adaptation and de-adaptation. Adaptation phases are shaded in blue in panels a and b. **c**. Walking on a treadmill with abrupt speed changes every 90 seconds^28^. Model qualitatively predicts experimentally observed step frequency changes. Transients have a fast and slow timescale, with the fast timescale change sometimes undershooting and sometimes overshooting the steady state (red and green detail). Without learning, with just the feedback controller (gray), the fast transient is preserved but the slow transient is replaced by a (noisy) constant. **d-i**. Walking with an exoskeleton that provides periodic propulsive impulses^48^. **d-ii**. Stride period converges to the perturbation period, implying entrainment. Different trajectories starting from different initial conditions are shown. **d-iii**. Perturbation phase converges to zero (heel-strike) in both model predictions and experiment. Different model trajectories show trials starting from different initial conditions. All quantities are non-dimensional. Source data are provided as a Source Data file.

The most popular experimental paradigm used to investigate human locomotor adaptation is walking on a split-belt treadmill^4,6,7,44^, which has two side-by-side belts that can be run at different speeds. Most humans have never experienced this novel situation. Humans adapt to walking on a split-belt treadmill on the timescale of seconds, minutes, and hours, exhibiting stereotypical changes in their walking motion^1,45,46^ and the model predicts these changes (Fig. 2a).

Specifically, within a few strides of split-belt walking, humans start walking with high negative step length asymmetry^4,44^ – that is, the step length onto the slow belt is longer than the step length onto the fast belt (see Fig. 2a and Supplementary Fig. 1e). This is the fastest timescale of adaptation, sometimes called ‘early adaptation.’ This negative step length asymmetry becomes close to zero over a few hundred strides (about ten minutes), and then becomes slightly positive with more time^7^. The model predictions have all these fast and slow transients both when minimizing just energy or energy plus a step length asymmetry (Fig. 2a and Fig. 8a). The model predicts an immediate initial increase in energy cost upon encountering the split-belt condition, which then reduces to a lower steady state gradually, as found in prior experiments^6^. When the split-belt condition is removed, the model predicts a fast-timescale transient to large positive step length asymmetry (a learning after-effect) and then a slow de-adaptation back to normal walking. The model predicts this de-adaptation to be faster than the adaptation, as found in experiments^4,7,44^ (Fig. 2a). The model also predicts that steady state is reached more quickly for step time asymmetry, and that the energy cost is more sensitive to step time asymmetry compared to step length asymmetry (Supplementary Fig. 3), as suggested by some prior experiments^6,47^.

**Figure 3.**
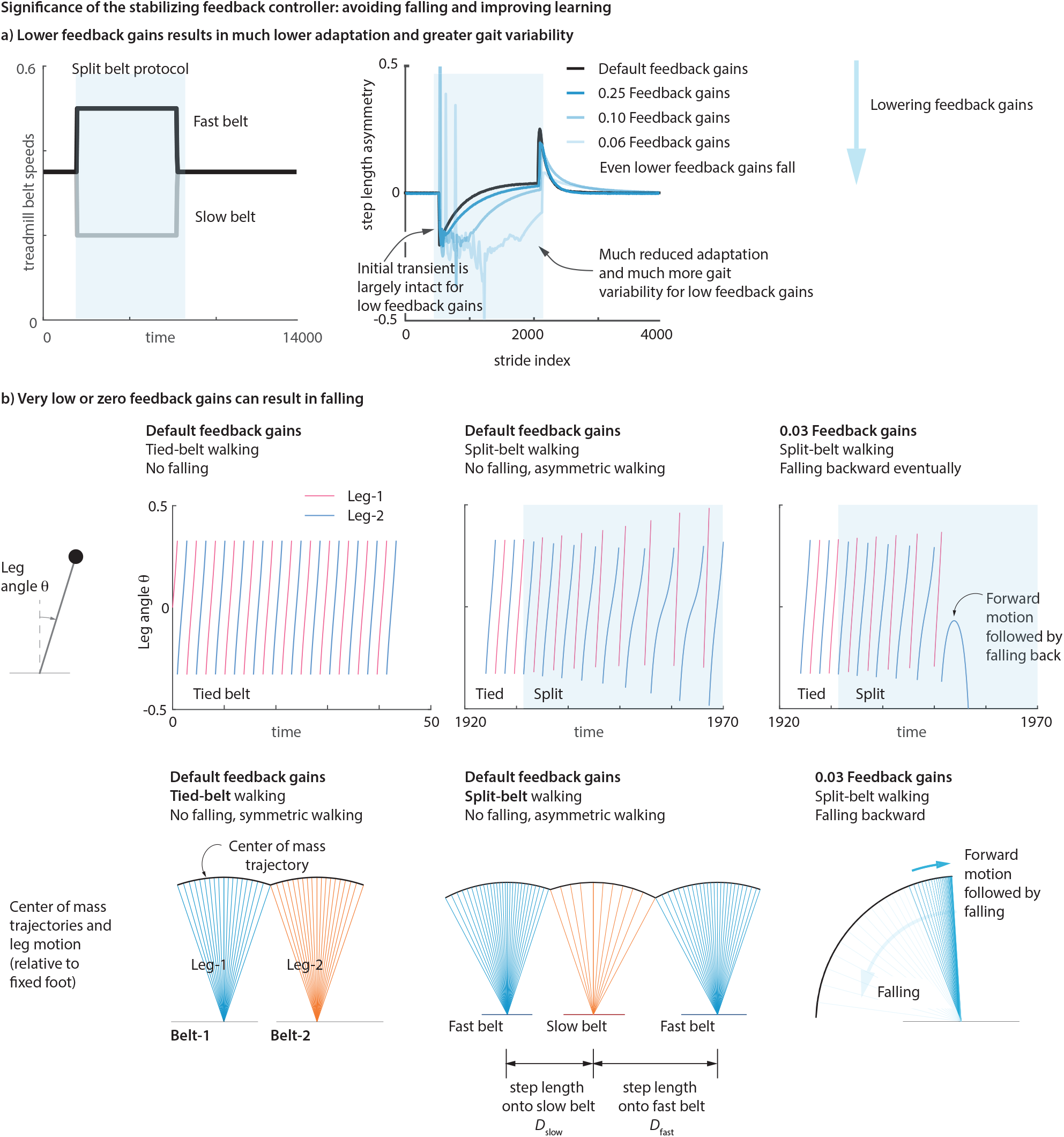
Significance of the stabilizing feedback controller: avoiding falling and improving learning. **a)** The default controller provides robust stability to the biped despite noise and environmental changes. Substantially lowering the feedback gains, all by the same factor, reduces the effective adaptation rate and increases gait variability. The sensory noise in these simulations is fixed across these feedback gain conditions and is applied to velocity feedback to the feedback controller. Adaptation phases are shaded in blue. **b)** Lowering the feedback gains even further results in falling of the biped upon introducing the split-belt perturbation. Three walking patterns are shown: normal tied-belt walking that has symmetric step lengths, split-belt walking with default feedback gains resulting in stable but asymmetric gait, and split-belt walking with much reduced feedback gains resulting in falling. In the bottom-most row, the center of mass trajectories for each stance phase are shown relative to the respective stance belt frame for visualization purposes (so that the split-belt trajectories for the different stance phases are with respect to different frames). Source data are provided as a Source Data file.

Human adaptation proceeds analogously when they are made to walk with an extra mass attached asymmetrically to just one ankle, as characterized by a prior experiment^5^. The model predicts the qualitative features of such adaptation, whether the performance objective is just energy or has an additional symmetry term (Fig. 2b). In both experiments^5^ and in our model, the walking gait becomes asymmetric in step lengths and then, during slow timescale adaptation, gradually tends toward symmetry; when the extra mass is removed, the asymmetry jumps to the opposite side, and then gradually de-adapts to normal walking.

The model predicts the step frequency changes while walking at varying speeds on a ‘tied-belt’ treadmill – which is just regular treadmill with one belt, or equivalently, a split-belt treadmill with equal belt speeds (Fig. 2c). In prior experiments^28^ in which the belt speed was changed every 90 seconds, humans quickly adapt their step frequency within 2 seconds and then slightly adjust their step frequency over a longer timescale — with the initial fast transient either overshooting or undershooting the ultimate steady state frequency slightly. In previous work^28^, the overshooting and undershooting transients required separate fits, whereas our model predicts both with the same framework.

The model captures empirical findings of how humans adapt to exoskeleton assistance. In some prior experiments^29,48^, humans were provided with time-periodic ankle torque impulses via a robotic exoskeleton (Fig. 2d). If the time period of these external impulses was close to the human stride period and the impulse magnitude was in the right range, the humans changed their stride frequency to entrain to this external impulse frequency, as predicted by the model (Fig. 2d-ii). Both model and experiment show entrainment that approximately aligns the external impulse with the transition from one step to the next (Fig. 2d-iii). The model can show entrainment whether the external impulse frequency is faster or slower than the stride frequency^29,48^, as found in prior experiment, while some prior models have shown that entrainment^29^ is possible for higher frequencies with just a feedback controller without learning. Rather than provide such time-periodic assistance, if the external assistive forces from the exoskeleton are a simple function of current body state (and not too noisy), the learner predicts successful adaptation toward the new optimum (Supplementary Fig. 4). We consider other such exoskeleton adaptation studies later in this manuscript (e.g., Fig. 4).

**Figure 4.**
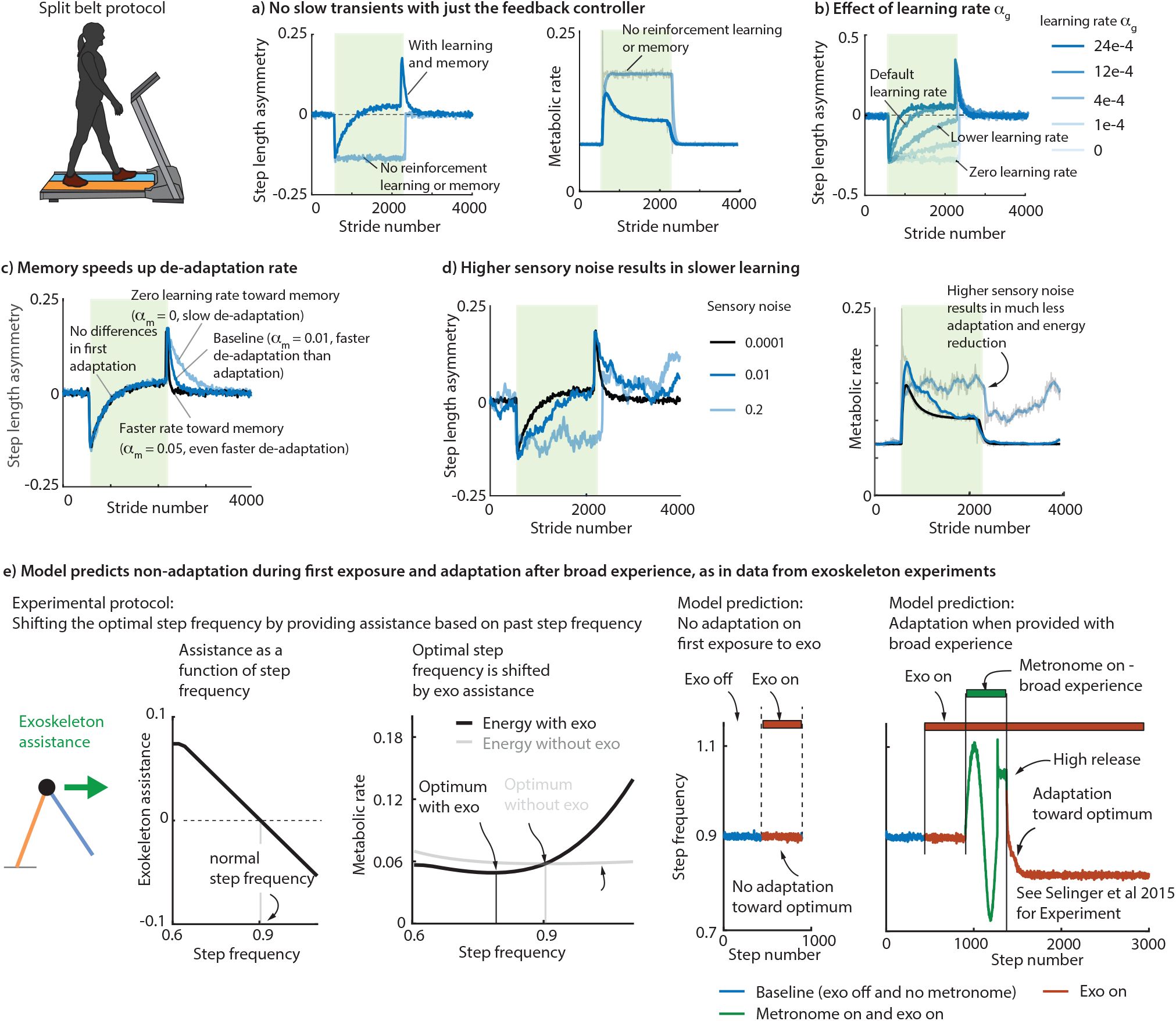
Effect of learning rate, memory and sensory noise: Model captures degraded learning, non-learning, and improving with guided experience. **a**. Stabilizing feedback controller alone only captures the fast learning transient. Addition of reinforcement learner is needed to capture the slow transients. **b**. Increasing the learning rate parameter speeds up learning (for a range of learning rates). **c**. Progress toward memory makes de-adaptation faster than adaptation. **d**. Increasing sensory noise degrades learning for fixed learning rate and fixed exploratory noise, resulting in less learning and less energy reduction. Split-belt adaptation phases are denoted by green shaded region in panels a-d. **e**. Model captures experimental phenomena^2,8^ wherein a human does not adapt to an exoskeleton that provides step-frequency-dependent assistance upon first encounter, but adapts toward the energy optimal frequency when provided with broad experience across a range of frequencies via a metronome-tracking condition. On the right two panels, blue indicates baseline condition without any assistance, red indicates exoskeleton assistance condition, and green indicates metronome-tracking condition in addition to exoskeleton assistance. In the rightmost panels, the ‘exo on’ condition (red) shows no adaptation before broad experience (green), but shows adaptation after the broad experience. All quantities are non-dimensional. Source data are provided as a Source Data file.

### Lesions in simulation identify modules responsible for the fast and slow adaptation transients

We can analyze which hypothesized modules in the model are responsible for explaining specific observations by the computational analog of ‘lesion experiments’: that is, turning off specific modules and noting what experimentally observed adaptation feature is degraded or lost. The following observations apply to all but the exoskeletal entrainment of the previous section, but we center the discussion on split-belt walking.

The fastest transient (early adaptation i.e., the initial response immediately upon experiencing the new condition) is entirely due to the default controller and the natural dynamics of the biped. Turning off both the reinforcement learner and the memory mechanism still results in the fast timescale initial response due to the stabilizing controller (Fig. 4a). Recent experiments partially corroborate this prediction, showing that providing gait stability through other means (e.g., handrail) affects this initial transient^49,50^, though such experiments may have changed other aspects of the gait than just stability.

Turning off the default stabilizing controller by setting all feedback gains to zero often makes the biped fall to the ground when the novel condition is initiated. Lowering the feedback gains to near zero results in falling or substantially degraded learning (Fig. 3a-b). Thus, the stabilizing controller is critical for effective locomotor adaptation. Further, this exercise of lowering the feedback gains closer to zero leaves a large fraction of the initial transients intact – showing that the feedforward component of this default controller is substantially responsible for the initial transient (Fig. 3a).

The slow adaptation transient when first exposed to the novel condition is due to the reinforcement learner improving performance. Turning off the reinforcement learner and the memory mechanism with zero learning rates results in the fast timescale initial response due to the stabilizing controller (Fig. 4a), but no slow timescale adaptation response. Thus, the stabilizing controller alone cannot explain the slow transients. Turning on the reinforcement learner results in the slow timescale adaptation response. Changing the learning rate for the reinforcement learner modulates the speed of this slow adaptation (Fig. 4b). In the first exposure to these novel situations, there is not yet any memory to call upon, and therefore, memory specific to the novel situation does not contribute to the first adaptation.

De-adapting to a familiar situation (equal belt speeds) after exposure to a novel situation will involve the use of stored memory of the familiar situation. Specifically, in split-belt walking, our model predicts that the de-adaptation will be faster than adaptation due to the use of stored motor memory of walking with tied-belts (Fig. 2a)^6,45^. Turning off this memory use, the de-adaptation is slower than adaptation (Fig. 4c). During first adaptation to a novel setting, the slow transients are governed by gradient descent, whereas during de-adaptation back to a familiar setting, the slow transients are sped up due to the summing of gradient descent and progress toward stored memory (Fig. 1d).

### Explaining savings, generalization, and anterograde non-interference

‘Savings’ refers to the faster re-learning of a task that has previously been experienced. In prior experimental work, such faster re-learning during a second split-belt adaptation experience was observed^1,51^, despite having a prolonged tied-belt period between the two adaptation periods (Fig. 5) — this intervening tied-belt period allows for full ‘washout’, complete de-adaptation in terms of observable variables. Here, our model qualitatively predicts such empirically observed savings (see Fig. 5 and Supplementary Table 1 for statistics). Such faster re-learning in the model is due to the motor memory mechanism, which stores how the controller changes under different situations. Motor memories are formed during first exposure to a novel condition, and then when exposed to this condition again, the re-learning is faster due to gradient descent and memory use acting synergistically (Fig. 1d). Because the motor memories are task-dependent, memories for split-belt adaptation do not decay entirely during tied-belt washout as the two tasks are non-overlapping. This persistent memory from the first exposure to split-belt walking results in the observed savings.

**Figure 5.**
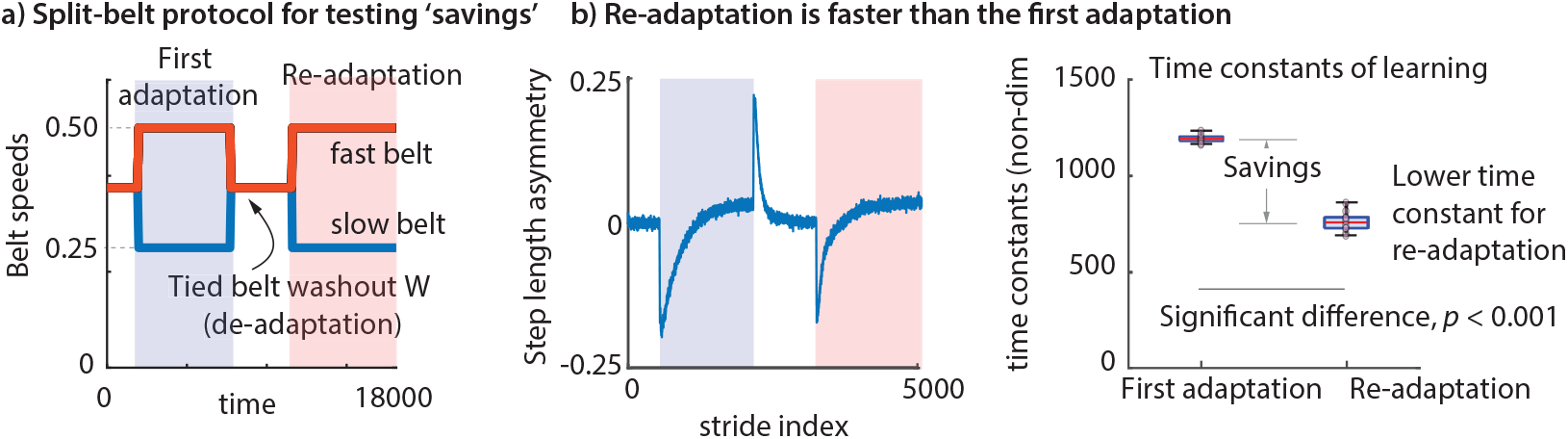
Savings. ‘Savings’ refers to the phenomenon that humans re-adapt faster to a condition (say, split-belt walking) if they have experienced the condition before, even if they have fully de-adapted to normal walking in the intervening time. a) A treadmill protocol with two split-belt adaptation periods with an intervening tied-belt washout de-adaptation phase (W) that brings all the externally observable state variables as well as the current controller back to baseline — but not the internal memory state, which remembers the past learning. First adaptation (blue shaded region) and re-adaptation (red shaded region) transients are shown. b) The model with memory predicts this experimentally observed savings phenomenon^51^. Step length asymmetry changes are faster during re-adaptation compared to the first adaptation. The re-adaptation has a smaller initial transient compared to the first adaptation. The time-constants of first adaptation and re-adaptation are computed by fitting a single exponential to the step length asymmetry transients, showing that re-adaptation has a faster time-constant. Source data are provided as a Source Data file. See Supplementary Table 1 for statistical details of comparisons. All box-plots show the median (red bar), 25-75% percentile (box) range (whiskers), and individual data points (pink circles).

‘Generalization’ is when adaptation under one task condition results in savings or faster adaptation for a different task condition. Humans exhibit generalization during locomotor adaptation and our model predicts this phenomenon (Supplementary Fig. 6a-b). Specifically, in one prior experiment^52^, humans exposed to a split-belt trial A showed savings for a split-belt trial B with a smaller speed difference between both belts than A. Thus, experience with task A sped up adaptation to task B, suggesting that humans generalized from A to B. Further, it was observed^52^ that such savings for task B from experiencing task A (with the larger belt-speed difference) was higher than the savings obtained if the first adaptation experience was with task B instead. Our model predicts both these generalization phenomena (Supplementary Fig. 6a-b) due to the motor memory being continuously parameterized with respect to continuous-valued task parameters (here, belt speeds), so that the controller for intermediate conditions are interpolated even if they are never directly encountered. Such generalization cannot be predicted by models in which memories are stored discretely without interpolation^2^.

**Figure 6.**
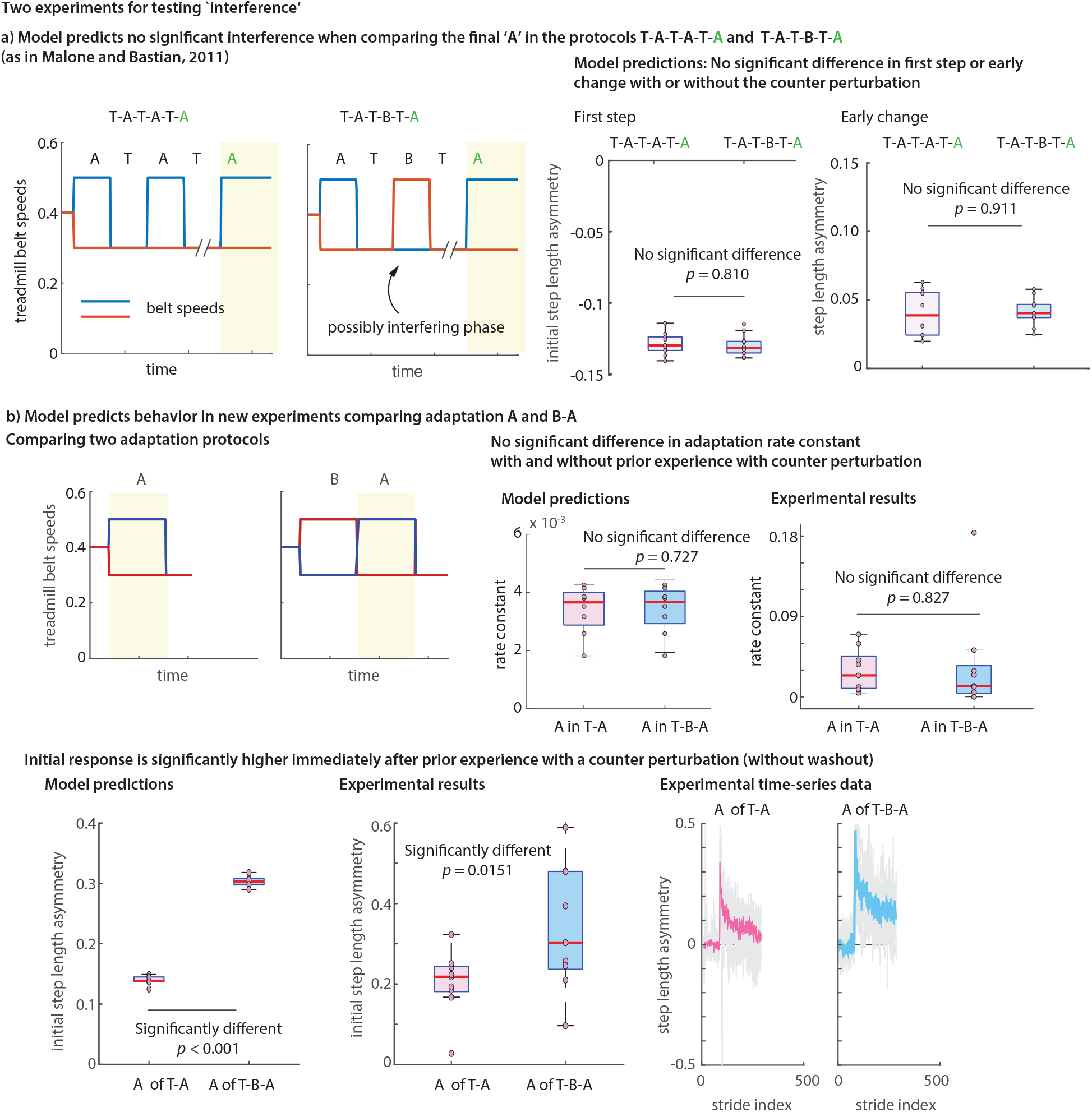
Interference. ‘Anterograde interference’ refers to the phenomenon where humans sometimes adapt slower to a condition A when they were previously exposed to the opposite condition B, that is, with the belt speed differences reversed between the two belts. a) We performed simulations of two split-belt adaptation protocols: first, T-A-T-A-T-A, alternating between tied-belt conditions T and the split-belt condition A, and second, T-A-T-B-T-A, where one of the A phases is replaced with the opposite condition B. We compare the adaptation between the two protocols during the final A phase (denoted as yellow shaded region). We find that the two protocols are not significantly different in their initial response to the perturbation or the early change in the step length asymmetry for the final adaptation period (yellow shaded region), as shown by Malone et al^51^. b) To test if this non-interference remains in the absence of washout, we performed prospective experiments in the absence of such a tied-belt washout phase: we compared protocols T-A with T-B-A with both simulations and human participant experiments. In experiments, we found that the initial step length asymmetry (first step of A) was significantly higher when B was present and the time constant of adaptation during A was not significantly different under the two conditions. This confirmed our model simulations, which predicted that the initial transients for A will be higher after B. The model also predicted no statistically significant difference in the adaptation rate constant in the presence of inter-participant variability of magnitude similar to that in the experiment. Box-plot shows median, 25-75% percentile and range. Source data are provided as a Source Data file. All box-plots show the median (red bar), 25-75% percentile (box) range (whiskers), and individual data points (pink circles). The time-series shows median as thick colored line and light gray lines are individual participant data overlaid. See Supplementary Table 1 for statistical details of comparisons.

‘Anterograde interference’ is when adapting to one task makes you worse at adapting to the ‘opposite’ task: opposite locomotor adaptation tasks could be split-belt walking tasks with belt speeds switched. Contrary to arm reaching adaptation studies where such anterograde interference is observed^41^, our model predicts that such interference need not happen in locomotion: that is, adapting to one perturbation need not make you worse at adapting to the ‘opposite’ perturbation if there is a sufficient tied-belt washout period between the two adaptation phases (Fig. 6a). This non-interference can be explained by the memory mechanism incorporating a function approximation, so that it can meaningfully extrapolate the learned controllers to the opposite perturbation as well. Such non-interference was indeed found in prior locomotor experiments^51^.

To further test the model’s predictions on how prior experience shapes adaptation, we performed prospective experiments here: we tested adaptation to two opposite split-belt tasks A and B without a washout period (see Fig. 6b), while prior experiments had a substantial washout period between the split-belt phases^51^. We found that the model predicted both the increased initial step length asymmetry transient due to the recent adaptation to the opposite task and the insignificant changes to adaptation time-constants (see Fig. 6b and Supplementary Table 1 for comparisons and statistics).

More generally, our model qualitatively captures effects of different split-belt adaptation protocols, for instance, capturing the time course of step length asymmetry when the split-belt phase is introduced gradually or abruptly, and whether these adaptation phases are short or extended^18,20^ (Supplementary Fig. 5). Having a longer duration adaptation phase in which the perturbation grows gradually may sometimes result in less savings than a shorter adaptation phase in which the perturbation began abruptly and remains constant (Supplementary Fig. 5). In previous work, an explicit memory of errors was used to explain some of these results^9^, but we have provided an alternative explanation via different model assumptions. In these cases (Supplementary Fig. 5), we found that the adaptation to different kinds of exposure to gradual and abrupt conditions can depend on protocol-specific parameters (e.g., duration of different phases, perturbation magnitude, learning rates); this suggests that one must be cautious of claiming general trends based on limited experiments.

The model predicts how the size and duration of perturbations affects adaptation^1,18,52^. In split-belt walking, both in model and in prior experiment^52^, being exposed to a larger belt-speed split results in larger initial transients and more positive final asymmetry (Supplementary Fig. 6c). Being exposed to a condition for a shorter period of time results in smaller savings than being exposed to the condition for longer^18^ (Supplementary Fig. 5).

### Degraded learning, non-learning, and making non-learners adapt via experience

The human motor system has sensory noise and motor noise that is not fully observable, and is thus distinct from intentional exploratory noise. The results presented thus far were obtained with low levels of sensorimotor noise. When the sensorimotor noise is less than a critical threshold, it preserves the qualitative results despite degrading the gradient approximation and thus degrading the effective learning rate (Fig. 4d). Large enough sensorimotor noise for fixed exploratory noise destroys the reinforcement learning entirely, resulting in no kinematic adaptation or energy reduction upon first exposure (Fig. 4d), potentially explaining why some populations with movement disorders may have impaired learning^53^.

Prior adaptation experiments involving exoskeleton assistance found that some humans were able to adapt spontaneously whereas others did not^2,8,27^. The non-spontaneous learners, when exposed to broad experience with a lower associated metabolic cost, were able to adapt toward the energy optimum^2,8^. In our model, both spontaneous learning and non-learning was possible depending on the size of sensorimotor noise: low noise resulted in spontaneous learning and high noise resulted in non-learning. As in experiment^2,8^, the model’s non-learners could be made to adapt toward a lower energy cost by giving them broad experience on the energy landscape, giving them experience of a lower energy cost to be stored in memory. In our model, this adaptation upon providing experience stems from motor memory formation and later memory use in addition to improving performance through gradient descent.

In addition to intrinsic sensorimotor noise, adaptation to external devices such as exoskeletons or treadmills could also be degraded by ‘device noise’. Our model predicts that split-belt adaptation can be degraded via such device noise when implemented as noisy belt speed fluctuations that are large enough (Fig. 7a). To test this model prediction prospectively, we performed human subject experiments and compared the post-adaptation after-effects of noise-free and noisy split-belt protocols. We found that participants had lower after-effects after the noisy adaptation condition, as predicted by the model; see Fig. 7a and Supplementary Table 1. This device-noise-based degradation may seem in conflict with earlier experiments by Torres-Oviedo and Bastian^20^, who compared adaptation in a split-belt protocol under noise-free and noisy belt speed conditions and found that the noisy version had higher adaptation as judged by the post-adaptation after-effects. However, our model also captures this improved adaptation due to different implementation of device noise in prior experiments^20^ by incorporating that specific protocol in the model (Fig. 7b), thus reconciling the seemingly conflicting findings. These results illustrate that the details of the noise pattern (e.g., magnitude and temporal correlations, see *Methods*) and the adaptation protocol used are important to determine the impact of device noise on adaptation, i.e., there are many ways to add device noise and some may enhance learning and others may degrade it.

**Figure 7.**
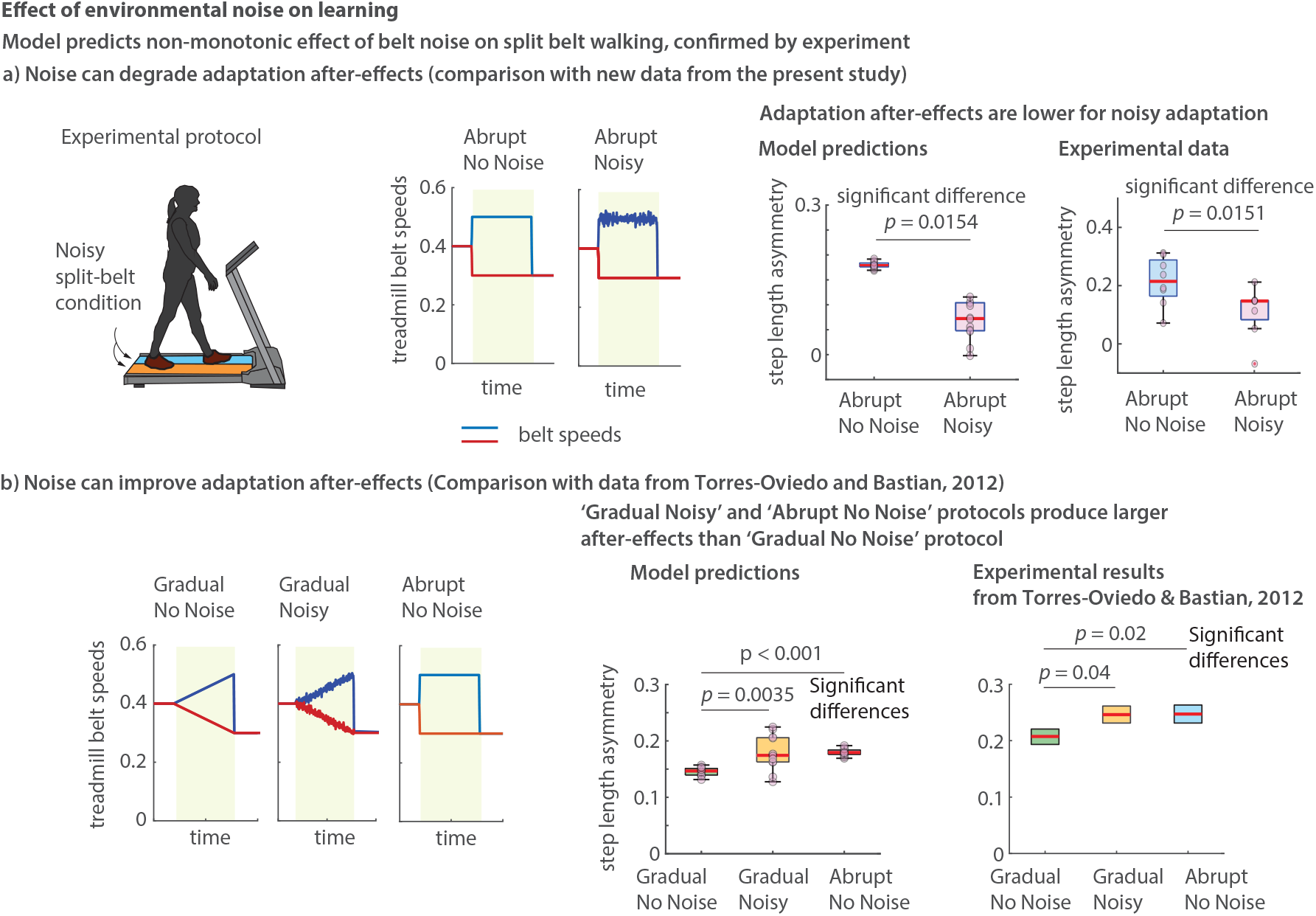
Split-belt learning can be degraded or enhanced depending on noise structure. a) Split-belt adaptation with and without belt-noise are compared. Adaptation phases are shaded in light green. The speed fluctuations in the noisy split-belt condition are continuous and piecewise linear and happen roughly every step. Model predicts that when the belt noise is high enough, adaptation can be degraded, as judged by lower post-adaptation after-effects. The post-adaptation after-effects shown are the initial step length asymmetry when the tied-belt condition starts after the split-belt adaptation. We performed model-guided prospective human experiments that confirmed these predictions: *p* value showing significant difference is from one-tailed t-test (unpaired, *t*(*d f*) =*−* 2.41(14), *p* = 0.0151). b) Torres-Oviedo and Bastian^20^ found that appropriately structured noise accompanied by gradual speed change can enhance adaptation, as measured by post-adaptation after-effects: box plot with these prior experimental results^20^ shows me n (red line) and standard error (box). Our model captures this behavior. All box-plots show the median (red bar), 25-75% percentile (box) range (whiskers), and individual data points (pink circles). See Supplementary Table 1 for full statistical details of comparisons. Source data are provided as a Source Data file.

**Figure 8.**
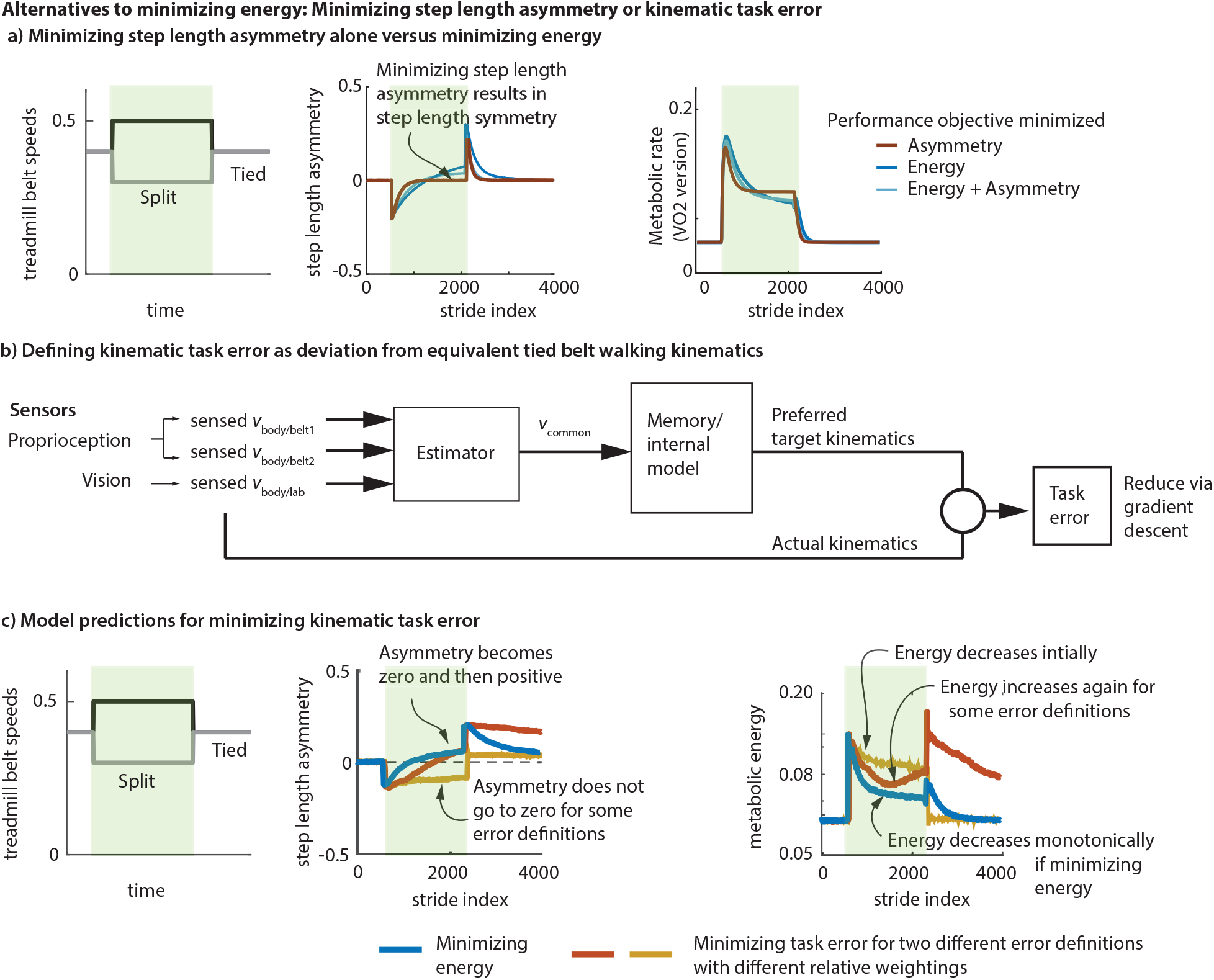
Alternatives to energy minimization: minimizing asymmetry or kinematic task error. **a)** As an alternative to an energy objective, minimizing step length asymmetry as the only objective during split-belt adaptation results in a perfectly symmetric gait in the model, which conflicts with the positive step length asymmetry found in experiment as well as when predicted by an energy minimization^45^. **b)** As another alternative to an energy objective, we formalize the minimization of kinematic task error as minimizing deviation from preferred walking kinematics defined for each speed. A single common belt speed is estimated by a state estimator using sensed body speeds relative to the lab and the belt (vision and proprioception). The task error is deviation from kinematics at that estimated speed under tied-belt conditions, drawn from memory. **c)** Model predictions for minimizing just task error without an energy objective; two different weightings are used for different components of the kinematic error (red and yellow). Energy minimization is shown for comparison (blue). For the task error predictions, one of either step length asymmetry or energy trends disagree with split belt adaptation experiments^45^: either the step length asymmetry stops well short of symmetry while decreasing energy somewhat (yellow), or the energy transients are not monotonically decreasing (red). Source data are provided as a Source Data file. Light green shaded region in all panels is the period of split-belt adaption.

**Figure 9.**
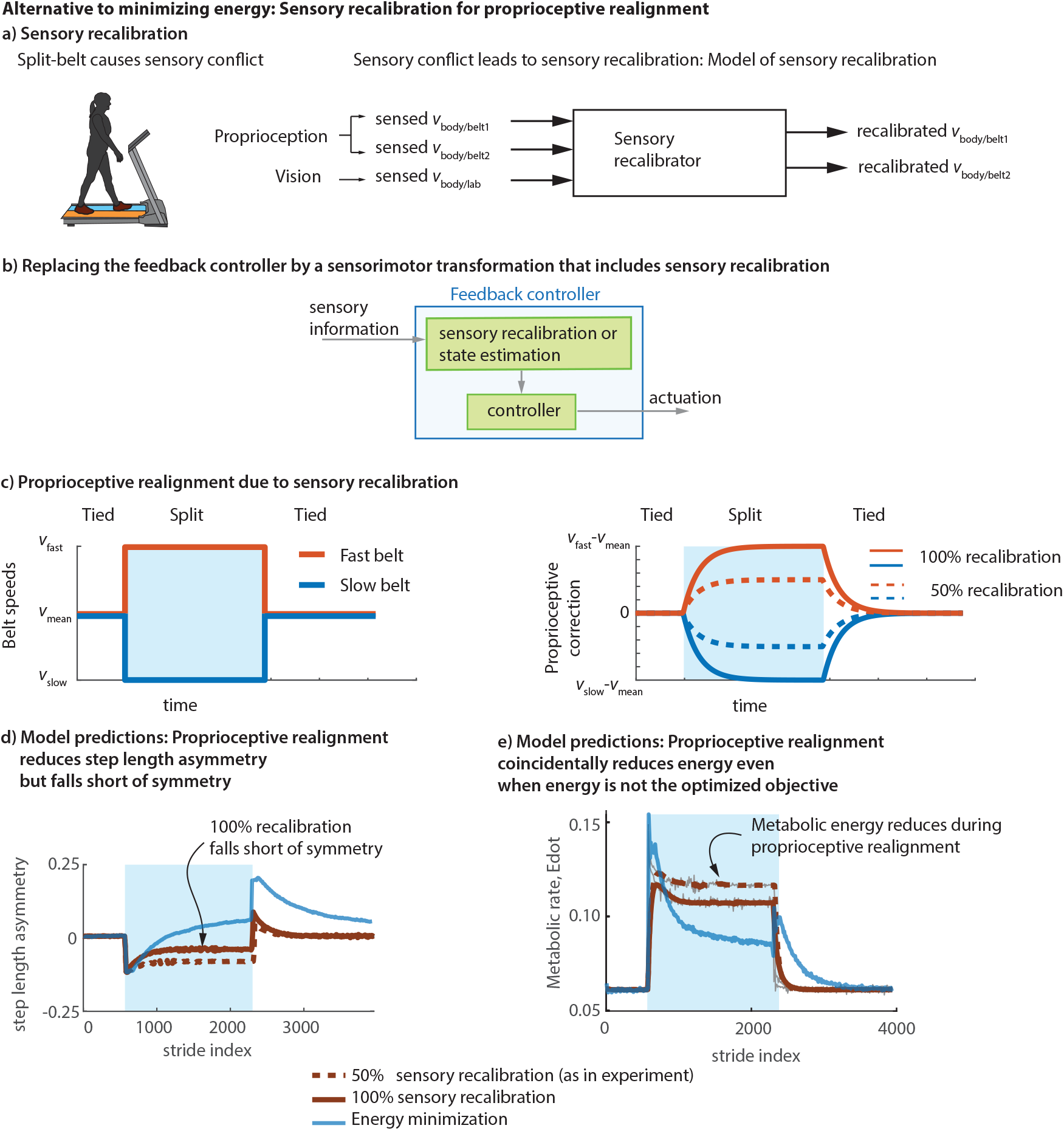
Alternative to energy minimization: Sensory recalibration via proprioceptive realignment. **a)** As an alternative to energy minimization, we considered sensory recalibration via proprioceptive realignment as a hypothesis for adaptation. This recalibration realigns the proprioception from the two legs to conform with the expectation that both legs usually are on the same surface with a common speed, rather than surfaces with different speeds. The proprioceptive recalibrator takes in the sensory information from proprioceptors of the two legs and vision and computes a recalibrated version of the proprioceptive information. **b)** The recalibrated sensory information is used by the the stabilizing controller, so the direct feedback controller of Fig. 1 is replaced by a more general sensorimotor transformation. **c)** Proprioceptive correction for each leg shown as a function of time for a particular split-belt protocol. This correction is subtracted from the initial proprioceptive estimate to obtain the recalibrated proprioception. The correction for the fast leg is positive and that for the slow leg is negative. 100% recalibration corresponds to completed proprioceptive realignment and 50% recalibration is close to that observed in experiment^42^. **d)** Using the recalibrated proprioception as feedback in the stabilizing controller results in the reduction of step length asymmetry, but even 100% recalibration does not result in symmetry or positive step length asymmetry. Thus, the model predicts that proprioceptive realignment cannot be fully responsible for split-belt adaptation. **e)** Proprioceptive realignment also reduces energy coincidentally even without energy being an explicit objective in this situation. Source data are provided as a Source Data file. Light blue shaded region in panels c-e is the period of split-belt adaption.

Aside from noise-based explanations, we provide one more potential cause for initial non-learning observed in some exoskeleton studies: delay between human action and exoskeleton response. Many exoskeleton adaptation experiments in which participants did not spontaneously adapt^2,8,27^ had an exoskeleton controller that provided assistance or resistance based on the participant’s previous walking step, resulting in a delay between action and energetic consequence. We showed that such delays can degrade or even stop gradient descent-based learning (Supplementary Fig. 7), making adaptation not obligatory. The gradient estimate is degraded due to poor credit assignment: when there is delay, the reinforcement learner in our model associates the effect with an incorrect cause, as the learner’s inductive bias assumes no such delays.

### Alternative to energy minimization: Comparing to minimizing asymmetry

To explain split-belt adaptation, researchers have treated the left-right asymmetry in step length as the error being corrected, fitting equations with one or two time constants to describe the observed decrease in this asymmetry^1,9,10^. Here, we examined what predictions our model makes if step length asymmetry is used in our optimization framework as the only performance objective, a variant of another study^24^ in which foot contact time symmetry was optimized. We find that minimizing asymmetry does not capture the slow timescale transients in either tied-belt or split-belt locomotion. First, for changing treadmill speeds during tied-belt locomotion^28^, minimizing asymmetry predicts the fast timescale changes in step frequency due to the default controller, but further slow timescale changes observed in experiment are not predicted by an asymmetry-minimizing objective alone. During split-belt adaptation, minimizing just step length asymmetry, our model predicts convergence to pure step length symmetry (Fig. 8a). This is in contrast to recent experiments which suggest eventual convergence to positive step length asymmetry^45^. In general, minimizing asymmetry is insufficient as the lone performance objective in an optimization framework, as perfect symmetry admits infinitely many locomotion patterns^54^ and does not result in isolated local minima required for stereotypy. Thus, minimizing asymmetry alone cannot predict the many steady state locomotor phenomena predicted by minimizing energy during normal locomotion^28,33,55^. As a corollary, when placed in any symmetric situation with a symmetric body (e.g., slopes or bilaterally symmetric exoskeletons), minimizing asymmetry will result in zero slow timescale adaptation of the controller even if the mechanical environment is changed substantially, in contrast to experimental findings^2,8,27^. While minimizing asymmetry alone does not explain diverse locomotor phenomena, minimizing a heuristically weighted combination of energy and asymmetry, with a small weight on the asymmetry, retains the qualitative predictions of minimizing energy, while sometimes allowing a better quantitative match (Fig. 2a-b and Fig. 8a). Future experiments could delineate the extent to which humans have symmetry as an explicit objective in addition to energy^56^, given that energy^54^ and other performance objectives such as proprioceptive realignment (as shown below) may also indirectly promote symmetry^22,42^.

### Alternative to energy minimization: Comparing to minimizing generalized task error

In low-dimensional adaptation tasks such as reaching with the arm to a target, the task error to be minimized is unambiguous; for instance, in reaching tasks with visuomotor rotation, the error is defined as angular distance to the reach target^9,12^. However, in higher-dimensional tasks like locomotion, analogous definitions of task error as deviation from desired body kinematics is not uniquely defined: for instance, the total task error could be defined as a weighted sum of the error from desired body states, with errors for different states weighted differently — but such a weighting would not be uniquely specified. Here, we considered a few such relative weightings and made model predictions for minimizing such kinematic task errors as the only performance objective (see *Methods* and Fig. 8b) via the exploration-driven gradient descent of Fig. 1b.

The resulting predictions were not entirely consistent with experiment. Different relative weightings resulted in distinct behaviors, all of which fell short of fully capturing the experimental findings: the weighting that results in eventual positive step length asymmetry, as seen in experiment, corresponded to energy increase in contrast to experiments, and on the other hand, the weighting that results in monotonic energy decrease has a steady state with substantial negative step length asymmetry, again in contrast to experiments (Fig. 8c). A purely kinematic performance objective was similarly found to not explain exoskeleton adaptation in prior experiments, where participants achieved entrainment to exoskeleton impulses^48^ or changed their walking frequency^8^ without plateauing at the unassisted walking kinematics.

### Alternative to performance optimization: Comparison with proprioceptive realignment

Proprioceptive realignment has been proposed as a potential mechanism accounting for the adaptation seen in split-belt locomotion^22,42^ and for arm reaching tasks with visuomotor perturbations^57,58^. Vasquez et al^42^ characterized the (proprioceptively) perceived speed of the legs after a split-belt adaptation, effectively finding that humans perceived the fast leg as being systematically slower than reality or the slow leg as faster than reality or both. A causal mechanism relating this sensory recalibration to locomotor adaptation has not previously been proposed, and a mathematical model could help establish if proprioceptive realignment could result in symmetry changes consistent with experiment.

We put forth a mathematical model of proprioceptive realignment via sensory recalibration using our framework, which enables linking body dynamics, sensory feedback (both proprioception and vision), and motor action. In our model, the two legs are expected to be on the same surface and proprioceptive deviations from this sensory prediction are perceived as an error to be corrected by recalibrating proprioception; while only proprioception is recalibrated, vision is used as a common sensory signal to estimate the proprioceptive conflict between the two legs (see *Methods* and Fig. 9a). This is a type of sensory prediction error^12,58^, as it is due to a difference between the sensory feedback and what the nervous system expects. This model results in recalibrated estimates of leg speeds such that on a split-belt treadmill, the fast leg feels slower and slow leg feels faster than reality, as in experiment^42^ (Fig. 9c), with the recalibration growing in time. The model produces no recalibration when walking on a tied-belt, as in experiment^42^. We incorporated this recalibrating proprioceptive sensing as feedback input to the stabilizing controller without changing other aspects of the default controller to predict what proprioceptive realignment alone can predict.

Proprioceptive realignment as implemented here falls short of explaining qualitative features of split-belt locomotor adaptation. Specifically, while the initial negative step length asymmetry produced by the default controller is decreased by proprioceptive realignment, the steady state of the adaptation still has substantial negative asymmetry (Fig. 9d), falling substantially short of experimentally observed symmetry^6,59^ and positive step length asymmetry^7,45^, which is predicted by energy optimization. Interestingly, the model shows coincidental metabolic energy decrease as a result of proprioceptive realignment (Fig. 9e), but this energy decrease is not accompanied by kinematic changes observed in experiment. Thus, while proprioceptive realignment could potentially be a partial cause of split-belt adaptation, it does not explain all the associated adaptation phenomena, as also suggested by recent experiments^60^. Beyond split-belt adaptation, proprioceptive realignment cannot explain how humans respond to tied-belt speed changes^28^, as experiments did not find significant proprioceptive realignment in the tied-belt condition^42^. Finally, proprioceptive realignment via interaction with vision, as implemented here, cannot explain adaptation to purely mechanical changes to the body or the environment such as an added mass or an exoskeleton.

### Interaction with explicit feedback

Our framework is meant to model implicit adaptation and learning, but can accommodate explicit adaptation mechanisms acting in parallel. One potential way to speed up locomotor adaptation is to provide explicit verbal instruction to the participant about the desired behavior or provide visual feedback on the error between desired and actual behavior^10^ (Fig. 10a). Indeed, providing visual feedback on step length asymmetry to participants on a split-belt treadmill and asking them to reduce this asymmetry hastened the progress toward symmetry — compared to adaptation without this feedback^10^. Removing this visual feedback partway through adaptation results in the increased symmetry being largely wiped out, so that the asymmetry goes back approximately to where it would have been without the explicit feedback. We were able to capture this phenomenon (Fig. 10c-d) by adding a separate module for explicit control that acts in parallel to the feedback controller in memory (Fig. 10a), as hypothesized in some prior work^10,23^. This demonstration is simply to show that the implicit learner of Fig. 1b can be readily modified to accommodate explicit mechanisms without degrading the implicit learner’s performance. This demonstration also shows that kinematic behavior changes due to explicit corrections need not, by themselves, be sufficient to modify implicit learning, as seen in experiments^10,21^.

**Figure 10.**
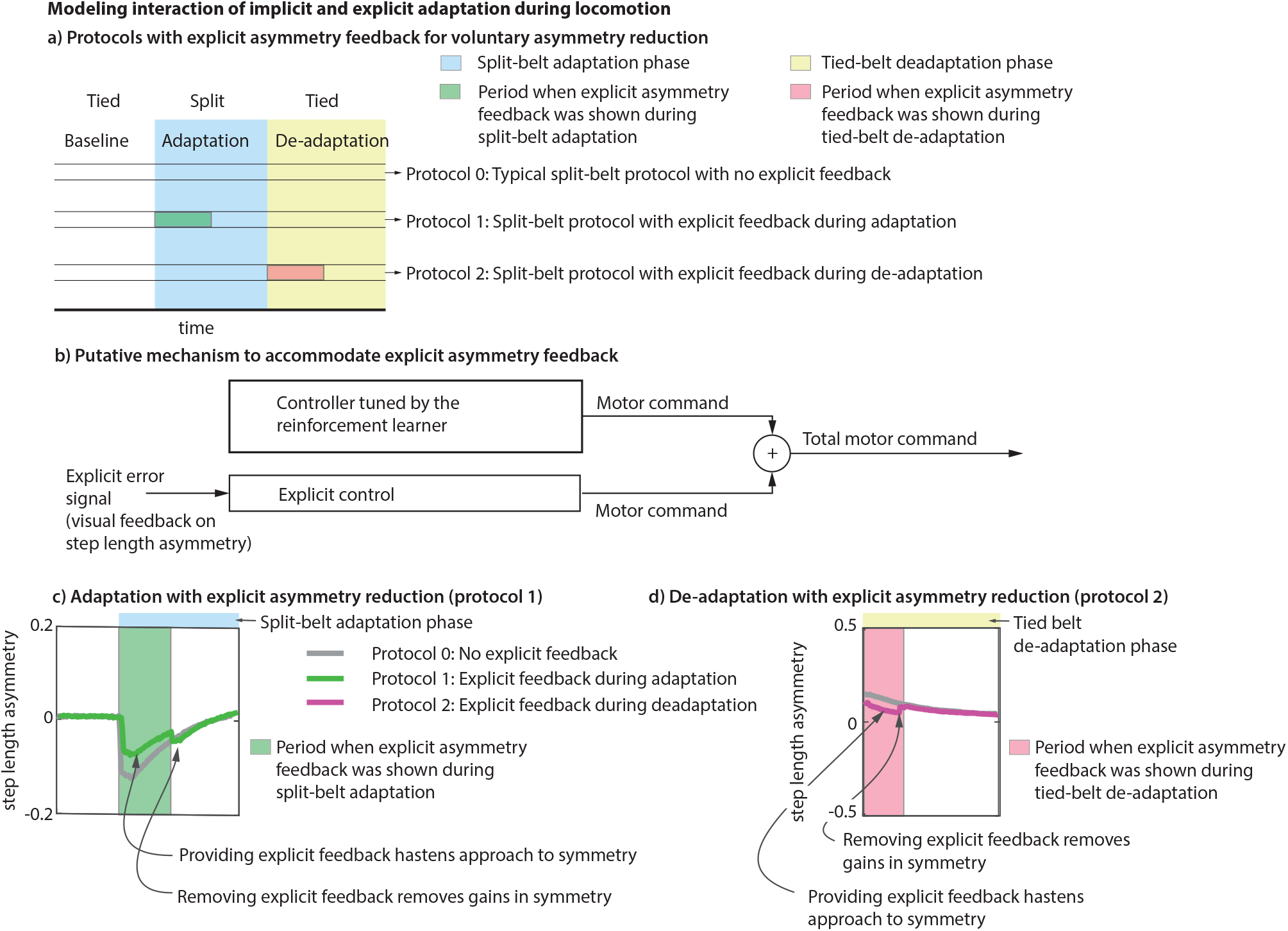
Interaction with explicit feedback. **a)** Three different split-belt protocols are compared. Protocol 0 which is a typical split-belt protocol with no explicit feedback about step length asymmetry. In protocols 1 and 2, participants are shown visual feedback of step length asymmetry on a screen and asked to reduce it explicitly, either partly through the adaptation phase (protocol 1) or partly through the de-adaptation phase (protocol 2). **b)** Explicit control is modeled as a module that adds to the nominal step length of the ‘implicit controller’ and this correction is proportional to the step length asymmetry on the previous step. The explicit and implicit modules are in parallel, and the implicit learner only knows about the motor command from the implicit feedback controller that it tunes. Having the explicit feedback improves progress to symmetry during **c)** adaptation and **d)** de-adaptation but this symmetry improvement is lost when the explicit feedback is removed, as found in experiment^10^. Source data are provided as a Source Data file.

## Discussion

We have presented a model for locomotor adaptation that captures observed experimental phenomena in ten different studies^3–8,18,20,28,48,51,52^, and predicts phenomena observed in two prospective experiments conducted in this study. Across these studies, our model captures adaptation transients in both the short timescale of seconds and long timescale of many tens of minutes. Our model also enabled us to compare different adaptation mechanisms, specifically energy optimization via reinforcement learning, proprioceptive realignment, and reducing sensory prediction error^22,42,58,61^, delineating how the hypotheses differ or coincide in their predictions and allowing testing through future prospective experiments. We have shown how humans could adapt to perturbations to the body or the environment, while walking stably and continuously without falling or stopping, as models of non-continuous episodic tasks such as arm reaching do not show how this is possible.

Predictive models of motor learning such as the one proposed here could be used to improve motor learning in the real world. We have made predictions about conditions that may degrade or accelerate learning consistent with prior experiments. Given this, future hypotheses for improving learning could be tested computationally within our modeling framework before testing via prospective experiments. We have tested the model by performing two such prospective experiments here, one for examining anterograde interference and another for the effect belt noise. Further such experiments may either provide further evidence supporting the model or information that could help improve the model. If the goal is to improve learning to use a device (such an exoskeleton or a treadmill), the device parameters and their sequencing can be optimized in simulation to reduce the time duration to learning steady state.

Our model suggests explanations for why humans may adapt reliably in some novel situations (for instance, during split-belt walking^4,6,20^) and not others (for instance, some exoskeleton studies^2,8,19,62^). One might wonder if common principles underlie such reliable adaptation in one class of devices and unreliable adaptation in another class, given that both devices are interacting with the same human motor control system. First, we note that both exoskeletons and split-belt treadmills share a core dynamical similarity: they are both mechanical devices that contact the body, applying forces and performing positive or negative work of different specifications^45^. Second, our results also suggest that the two classes of devices are not fundamentally different with respect to motor learning, but that the differences may be due to dynamical properties of treadmills versus current exoskeletons and their controllers. Our model suggests ways in which we can make participants less reliable learners on split-belt treadmills and reliable learners on exoskeletons and prostheses. Specifically, our model predicts that split-belt adaptation can be degraded by noisy belt speed variations (Fig. 7) — which we confirmed with our prospective experiment. We also noted that many exoskeleton studies that did not show obligatory adaptation involved exoskeleton controllers that had a one step delay between human action and the device response^2,8,19^. We showed that gradient descent can be substantially degraded or entirely stopped in the presence of such delays (Supplementary Fig. 7), whereas there can be reliable learning in exoskeletons with no delay or noise (Supplementary Fig. 4); this prediction can be tested by systematically manipulating the device delay in future experiments. In summary, we suggest that humans may exhibit better adaptation to exoskeletons if the device has low noise, has simple consistent dynamics from step to step, and does not have substantial delay between human action and device response.

A corollary to the prediction that lowering device noise improves learning reliability is that increasing baseline human exploratory variability compared to unresolved sensorimotor or device noise may improve learning reliability. It is an open question whether baseline exploration as used by the nervous system in implicit learning can be manipulated by an experimenter via purely external means (that is, via sensory or mechanical perturbations or other biofeedback) — in a manner that results in more reliable learning. One study that increased variability externally did not find better learning^63^, while another study increased learning^20^: our model was able to recapitulate the increased learning in the latter study. Another study performed a manipulation that increased both variability and learning^37^. It is unclear if this increased variability specifically corresponds to increased exploration because both studies changed the sensory or the mechanical environment, which could have increased variability by increasing unresolved sensorimotor noise. Further, according to our model, such increased variability comes with a higher energetic cost at steady state^2^ as well as potentially higher fall risk, so future work could use our model in concert with targeted experiments to delineate how humans trade-off these competing objectives of exploration, energy, and stability.

Our model naturally predicts the various qualitative features of short timescale and long timescale responses to perturbations without fitting to the adaptation phenomena being explained. This is in contrast to the single rate or dual rate or memory of errors models of adaptation^9,18,46^, which when applied to locomotor adaptation without including bipedal dynamics and control, do require fits or specific assumptions to capture the direction of both the slow and the fast timescale transients. Here, we predict the short timescale response to sudden perturbations as simply the response of the default stabilizing controller to those perturbations, and this prediction obtains the correct direction or sign of the response without fits to the data it tries to predict. For instance, our model naturally predicts that the immediate transient upon a split-belt perturbation or a leg mass addition is negative step length asymmetry (Fig. 2-4). Similarly, we have shown that a substantial part of slow timescale motor adaptation can be predicted by performance optimization, with energy consumption as the performance objective. This model obtains the correct direction of the slow adaptation without any fits to the adaptation data. In contrast, in the traditional dual rate or memory of errors adaptation models^9,11,18^, the direction of slow adaptation is toward zeroing the error and, therefore, is dictated by how error is defined. Thus, while descriptive models^9,11,18^ may be fit to short and long timescale transients in some locomotor adaptation experiments, they do not make predictions of the transients from more primitive assumptions. In addition, we have shown that some common ways of defining error, when coupled with locomotor dynamics, may result in predictions that disagree with experiments.

Our accounting of savings and memory is complementary to previous work that have addressed savings or other related phenomena via memory mechanisms centering on context inference for error-based learning or for performance improvement^2,41,46,64^. These previous works did not consider the interaction of performance improvement and stabilizing control in a complex task such as locomotion^41,46,64^, as here, or when considering locomotion, did not consider locomotor dynamics and control^2^. Our memory model is also different from models that adapt the ‘error sensitivity’ (learning rate) of adaptation via a memory of sensory errors^9^, which can capture savings in the form of faster adaptation rates, but is similar to other linear time-invariant state-space models^46^ in that neither model can capture savings in rate after a complete washout^65^.

We have argued that predicting human locomotor adaptation phenomena may require the following functional components: a stabilizing controller, an optimizing reinforcement learner, a gradient estimator, a memory mechanism, and possibly a module that reduces sensory errors. Like all mathematical models of complex phenomena (famously in string theory^66^), there may be multiple realizability: the same architectural hypothesis can be expressed in different terms, grouping some components together, dividing components into their sub-components, or have different realizations of similar function. No matter this multiple realizability, we have shown that a necessary feature of locomotor adaptation is exploration in the neighborhood of a stabilizing controller. Further, the framework implies the existence of a hierarchical separation of timescales of the model components^67^. Specifically, the step-to-step stabilizing controller has the fastest timescale, matching the timescale of the bipedal dynamics to prevent falling; the timescale of gradient estimation must be slower than the step-to-step dynamics so that the estimated gradient is reliable; finally, the timescale of the local reinforcement learner must be slower than the gradient estimator, so that the learner does not change the parameters too quickly for the gradient estimate to be reliable. Human motor learning proceeds over multiple timescales^1,46^, and our approach thus provides a natural functional account of the hierarchy of these timescales from the necessity of stable learning^67^.

Our model of locomotor adaptation is hierarchical and modular. Evidence for the hypothesis of hierarchical and modular motor control goes back to hundred year old experiments in which decerebrate cats produced coordinated repetitive movements but not goal-directed movements^68^. It is thought that fast timescale motor responses may be mediated in part by spinal circuits while longer-timescale control, adaptation, and context-dependent responses may be achieved by interaction of the cerebellum and motor-related areas of the cerebrum^3,24,69,70^. In our model, we have separated the fast timescale stabilizing controller and the slow timescale adaptation mechanisms into distinct interacting modules, so that damage to just the slow timescale adaptation module in the model could still preserve the fast timescale stabilizing response. Such preservation of fast timescale response to treadmill speed changes with degraded slow adaptation to a split-belt condition was found in participants with cerebellar damage^71,72^. Indeed, such studies have established that one locus of such slow timescale motor adaptation, especially involving sensory recalibrations and internal model change, is the cerebellum^3,11,12,71–74^. Thus, while our model is meant to be at the Marr level 1 and 2 (computational and algorithmic levels)^75^, it could inform interpretation of data on neural underpinnings. Conversely, neural data may allow us to fine-tune our model architecture: for instance, modules in the model may contain sub-modules responsible for distinct aspects of behavior which may be neurally dissociable (e.g., spatial and temporal slow adaptation^44,74^). Some studies have suggested preservation of ‘reinforcement learning’ despite cerebellar ataxia^11,13,76^, but such studies examined learning from explicit visual or auditory feedback, which is distinct from the implicit reinforcement learning we have proposed for energy optimization.

Human motor control strategies in highly practiced and learned tasks tend to approximate optimal controllers^77^, and here we have provided an account for how humans gradually learn such optimal controllers in a novel environment. A related learning paradigm is that the nervous system gradually learns an inverse model of the task dynamics from unsuccessful trials, and then uses the inverse model to achieve the task^78^. However, such inversion does not have a unique solution in high-dimensional tasks such as locomotion: human bodies have infinitely many ways to solve a movement task^26^ and thus must usually optimize another performance objective to obtain a unique solution. Here, gradient descent of the stabilizing controller implicitly accomplishes both the inversion and the optimization, as the resulting controller performs the task while optimizing performance.

Our model demonstrates that a local exploration-based search strategy and a simple linear controller structure is sufficient to describe the continuous adaptation of locomotion by human adults to changes to their body and their environment, starting from a known default stabilizing controller, learned under normal conditions. Our approach may lend itself to comparison with the recently popularized framework of deep reinforcement learning^79–81^, which use more expressive controller approximations (deep neural networks) with orders of magnitude more parameters. These methods do not assume initialization with a default controller but instead employ highly exploratory search involving thousands of discrete walking episodes, often involve falling and resetting the initial condition at the end of each episode. Thus, these learning methods operate in a different regime from our model and are not aimed at explaining gradual human locomotor adaptation.

Most learning requires trial and error, but attempting to improve locomotion via simple trial and error without a stabilizing controller as an inductive bias can result in falling or other learning instabilities. The stabilizing controller in our model allows safe exploration and adaptation, and turning off the stabilizing feedback while the gait is adapted results in falls or at least substantially degrades learning (Fig. 3). This shows that what control policy the learning acts on determines the effectiveness and safety of the adaptation. We also found that a number of alternative choices can result in falling: prioritizing energy optimization over the near future rather than over a longer time-horizon, too high a learning rate, and updating the gradient estimate too quickly. We have posited the use of exploratory variability for reinforcement learning or optimization, as also suggested in a few studies^2,11,13,38^, including experimental evidence for the role of exploration in improving error-based learning^38^. It was not known how such exploration could be implemented to adapt while walking continuously, without ignoring the locomotor dynamics, stability, and the continuous nature of locomotion (i.e., not treating each step as an independent episode). Indeed, using simple trial and error to perform optimization, for instance, using an exploration-driven search depending on just the previous step^2,13^, works for episodic arm reaching but will result in falling or non-learning for walking with continuous locomotor dynamics. Thus, here, we have put forth a framework for predicting how humans adapt their walking to different conditions while continuing to be stable.

We have tested our model against a wide variety of adaptation studies, providing broad empirical support for the model’s predictive ability. Future work can involve the design of targeted experiments to test the different components of this model (e.g., performance objective, adaptation algorithm)^82^, as these components contain heretofore untested assumptions about locomotor adaptation. Here, we have compared the predictive ability of performance objectives such as energy, symmetry, and sensory prediction error, determining what each can predict when acting alone. Future experiments can systematically manipulate the energy landscape, sensory feedback (e.g., vision), and unforeseen perturbations during adaptation to delineate how these performance objectives are traded off by the human nervous system^83^ — our model, which allows these adaptation mechanisms to act simultaneously, can provide a framework for interpreting such experiments. Here, we have shown the sufficiency of exploration-based gradient estimation and gradient descent with a fixed learning rate in predicting diverse adaptation phenomena. Future experiments can compare the predictions of gradient descent versus alternative descent or adaptation algorithms (e.g., gradient descent with momentum^39^ or learning rate adaptation^9^) in long timescale trials that either have gradually time-varying conditions or alternate between different conditions at various switching frequencies. Such prospective experiments would allow us to characterize the relation between the adaptation direction in experiment and the model-predicted gradient directions, thus helping to modify the model to capture a broader range of experiments. Future work can also test the generality of our framework to other motor adaptation tasks^41,77,84^, including the model’s ability to explain savings, generalization, interference, non-learning, and other important phenomena; this application of our model to other motor tasks will require appropriate modifications to the dynamical model and the default controller.

Our focus has been on capturing qualitative phenomena and we did not obtain a quantitative fit by minimizing the error between model predictions and experiment. Consistent with this preference, we used a simple biped model with simplified actuation, sensing, and default controller structure, which was sufficient for broad qualitative predictions but may limit ability to produce detailed quantitative predictions; indeed, model simplicity may be a sound reason to not seek quantitative fits. While we have captured a wide variety of experimental phenomena from diverse labs, future work could use a higher dimensional musculoskeletal^56^ and sensorimotor model and test it against other prior experimental data not considered here^60,85–87^ in addition to the aforementioned prospective experiments. In these future studies, we would seek quantitative fits to many aspects of the experimentally observed adaptation behavior (e.g., detailed kinematics, kinetics, energetics, variability), not just the time course of one or two variables (as is typical) and without experiment-specific parameter tuning.

Model-based predictions of locomotor adaptation, such as enabled here, have potential applications to improving human-machine interactions including robotic prostheses and exoskeletons, making such devices intrinsically more learnable or devising protocols for accelerating their learning^56,86,88^. Comparisons of learning in healthy and impaired human populations^53^ using our modeling framework provides a means of identifying how distinct hypothesized modules of locomotor adaptation may be affected, potentially informing targeted rehabilitation.

## Methods

In this Methods section, we first describe the mathematical structure of each component of our modular and hierarchical locomotor adaptation model (Fig. 1), how the components interact, and how this framework is applied to each task setting; the human experiments are described at the end. Human participant research reported herein was approved by the Ohio State University Institutional Review Board and all participants provided informed consent.

### Stabilizing feedback controller

The mathematical biped model, approximating the human walker, is controlled on a step-to-step basis by a feedback controller. The biped model and the stabilizing feedback controller^16,26,31,89^ are described in greater detail later in this *Methods* section (see also *Supplementary Methods*). Here, we describe the general structure of the feedback controller necessary to understand our modeling framework. The feedback controller is a function that relates the control variables *u* (e.g., forces and torques) to the the state variables *s* (positions and velocities). Here, the state *s* is a vector with as many elements as there are state variables (*n*_state_ elements) and analogously *u* is a vector with *n*_control_ elements. The control variables have nominal values *u*_nominal_, sometimes referred to as a ‘feedforward’ term, which the biped uses in the absence of any perturbations at steady state. Analogously, the state variables also have nominal values *s*_nominal_ in the absence of any external perturbations. Then, on step *j*, the control variables *u*_*j*_ are assumed to be related to the state *s*_*j*_ by the linear equation:

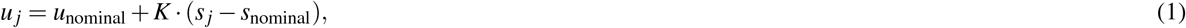

where *K* is an *n*_control_ × *n*_state_ matrix of feedback gains. This equation 1 is equivalent to the simpler linear expression *u*_*j*_ = *a* + *K·s*_*j*_, which has fewer parameters because the two vector variables *u*_nominal_ and *s*_nominal_ in equation 1 are replaced by the one vector variable *a* = *u*_nominal_ *K·s*_nominal_. This vector *a* may be considered the full ‘feedforward component’ of the controller, in that it contains all terms that do not directly depend on current state. We use the version including *u*_nominal_ and *s*_nominal_ in equation 1, in order to demonstrate the learner’s ability to automatically ignore redundant parameters. The linearity of equation 1 is a simplifying assumption, justified by the ability of linear controllers to explain human step to step locomotor control^16,25,31,89^ and its sufficiency for the adaptation phenomena explained by the framework here. The framework itself does not rely on this assumption of linearity.

### Local reinforcement learning for performance improvement

When faced with a novel situation, the reinforcement learner changes the parameters of the stabilizing controller to make progress toward a defined objective, expressed as minimizing a scalar objective function or performance objective *J* evaluated over each stride. The learnable parameters *p* characterizing the stabilizing controller include *u*_nominal_, *K*, and *s*_nominal_, i.e. the nominal control and state values as well as the feedback gains. In this study, we only allow the nominal values *p* = [*u*_nominal_; *s*_nominal_] to change during learning. This is because there is a one-to-one mapping between these nominal or feedforward terms and the overall gait kinematic changes we are trying to predict, so allowing the nominal values to change gives the model sufficient flexibility to produce different kinematics. We keep the feedback gains *K* fixed, as the primary role of the feedback term is to keep the system stable despite fast timescale perturbations away from the current gait pattern. Given the robustness of the controller to substantial perturbations^16,17^, this stabilizing role is satisfied by fixed feedback gains *K*. Indeed, as assumed, we find that changing them is not necessary for the major phenomena discussed herein; allowing just the feedforward term to change^3^ is sufficient (e.g., Fig. 2a-d). Allowing the feedback gains *K* to change may be necessary for even more stability-challenging perturbations, where the robustness of the default controller no longer is sufficient — such changes to *K* can be accomplished with the same framework but would require incorporating the locomotor task constraints explicitly into the performance objective (e.g., not drifting off a finite treadmill, not falling, traveling a certain distance), as otherwise the feedback gains may be chosen in a manner that makes the walker unstable. During learning, we allow the *u*_nominal_ and *s*_nominal_ to change independently for the left and right steps, enabling adaptation to asymmetric conditions.

Fixing the overall structure of the controller during learning to that in equation 1 makes this initial controller structure an inductive bias for learning; that is, it constrains exploration-based learning both by providing an initial condition and restricting the space of controllers explored.

On each stride *i* (every two steps), we denote *p*_*i*_ to be the current best estimate of the controller parameters. We posit that before encountering a novel condition, the body uses the previously learned controller for normal walking, which we have characterized using data from normal walking^16,31,89^. We term this the ‘default controller’ with parameters *p*_default_, so that on the first stride, the parameters are *p*_1_ = *p*_default_. Given the controller parameters *p*_*i*_ on stride *i*, the reinforcement learner chooses the controller parameters for the next stride *p*_*i*+1_ as the sum of two terms: the old controller parameters from the previous stride (*p*_*i*_) and a small change along the negative of the gradient estimate of the performance objective:

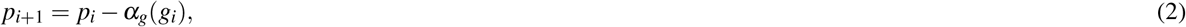

where *g*_*i*_ is the current gradient estimate on the *i*^th^ stride (see equation 7 for how it is estimated) and *α*_*g*_ is a scalar learning rate for the gradient descent. Rather than executing the next stride using this new *p*_*i*+1_, we posit that the nervous system uses a perturbed version 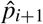:

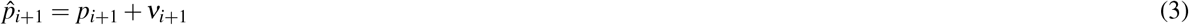

where *ν*_*i*+1_ is an exploratory motor noise term, assumed to be multivariate Gaussian noise with standard deviation *σ*, uncorrelated across time. We posit that the exploratory noise *ν* is intentionally generated by the nervous system, allowing it to estimate the local gradient of the performance objective *J* with respect to the parameters *p*, and more generally, build a local internal model of the system, serving as persistent excitation in the parlance of system identification^90,91^. This exploration-based estimation of the gradient is in contrast with other simulation-based ways of estimating the gradient, for instance, algorithmic or automatic differentiation, also called backpropagation^24,79^. In addition to this exploratory motor noise, there may be additional unavoidable sensory and motor noise that the nervous system cannot resolve, which we consider later separately^92,93^. The proposed reinforcement learning procedure directly updates the parameters of the control policy via gradient descent, so it may be considered a variant of policy gradient reinforcement learning, where the gradient is estimated as below entirely from exploratory steps^94^. Because the gradient is updated from limited and noisy data (see below), it is a stochastic gradient descent on the control policy. We term this learning ‘local’ because of its reliance on the information obtained via local exploration in the neighborhood of the controller to make gradual progress toward the optimum. In this formulation, the learnable parameters *p* are changed every stride, so that the effect of the left and the right step control on the performance objective can be experienced separately before being incorporated into the parameter change. This assumption of once-a-stride parameter change is not essential; the learning framework can be used with continuous phase-dependent control^25,31^ with more frequent or continuous updates of control parameters.

### Asymptotic gradient estimate

Estimating the gradient of the performance objective *J* with respect to the parameters *p* is equivalent to building a local linear model relating changes in parameters *p* to changes in performance *J*. This can be understood by noting that a local linear model is the same as a first order Taylor series, and the gradient ∇_*x*_ *f* of a function *f* (*x*) about *x*_0_ appears as the coefficient of the variable *x* in this first order Taylor series as follows:

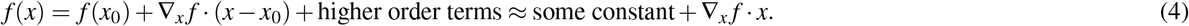

The performance *J* on a given stride will not only depend on the controller parameters *p*, but the entire system trajectory, which is uniquely determined by the initial system state and the subsequent control actions given by *p*. So, we posit a linear model that includes dependence on both *s*_*i*_ and *p*_*i*_. On stride *i*, if the initial state is *s*_*i*_, the parameters are *p*_*i*_, and the performance over that stride is *J*_*i*_, a linear model relating these quantities is given by:

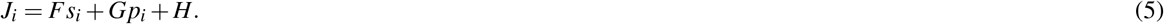

Here, coefficient matrix *G* is the gradient of the performance *J*_*i*_ on the current stride with respect to the learning parameters *p*_*i*_ and the coefficient matrix *F* is the gradient with respect to initial state *s*_*i*_. Building a linear model of *J*_*i*_ with respect to only *p*_*i*_, ignoring the dependence on state *s*_*i*_ can lead to incorrect gradient estimates and unstable learning.

Performing gradient descent using the matrix *G* as the gradient is equivalent to reducing the performance of a single stride *J*_*i*_, without considering the long-term implications. Minimizing just the single-stride performance *J*_*i*_ may result in unrealistic optima for some performance objectives: turning off the actuators and falling may be optimal when only minimizing energy over one step. So, for non-transient tasks such as steady walking, we hypothesize that the human prioritizes the long term or steady state performance *J*_∞_ = lim_*i→*∞_ *J*_*i*_. This asymptotic or long-horizon performance averages over the noise on any one stride.

To estimate the gradient with respect to long term performance, the nervous system needs to be able to predict the future. Thus, to predict the long-term consequences of the parameters *p*_*i*_, we posit that the nervous system maintains an internal forward model of the dynamics, that is, how the initial state *s*_*i*_ and the parameters *p*_*i*_ for a stride affects the state at the end of the stride, equal to the initial state for the next stride *s*_*i*+1_. This internal model of the dynamics is also assumed to be linear for simplicity:

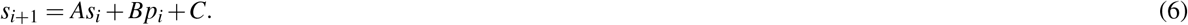

Given such an internal model of the dynamics, the nervous system can estimate the future consequences of parameter changes to the steady state (by effectively simulating the internal model to steady state) and thus infer the relevant gradient of *J*_∞_, given by:

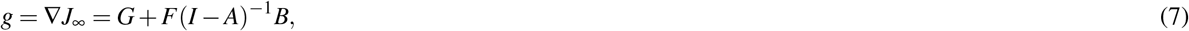

where the first term *G* gives the gradient of the short term energy cost over one step, while the second term corrects for the fact that the steady state value of *s* will be different from the current initial state *s*_*i*_. This internal model framework also allows the nervous system to minimize performance over an intermediate time horizon by computing and using the gradient of the mean energy cost over the next few strides. We found that minimizing expected performance over just one or two strides into the future can result in unstable learning for energy optimization. In conventional reinforcement learning^94^, a discount factor 0 *< γ <* 1 is used to modify the function minimized to 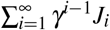, which prioritizes near term performance and down-weights performance in the future. We did not use such a discount factor here, but using *γ≈* 1 is analogous to the asymptotic limit we have chosen, and *γ* much less than one will give results similar to optimizing over just the next few strides.

We update the matrices *A, B, C, F, G, H* on each stride by estimating the linear model via ordinary least squares to best fit the state, the action, and the performance (*s*_*i*_, *p*_*i*_, *J*_*i*_) over a finite number of previous steps. We used a rolling estimate over 30 steps for all the results presented herein. This gradient estimator needs to have the property that relatively prioritizes recent history^90^, as otherwise, the gradient cannot adapt to novel locomotor situations in a timely manner. Using a finite history allows rapid adaptation to sudden changes. Also, we use a linear internal model though the full biped dynamics are nonlinear; a linear internal model is sufficient when adaptation is gradual and the model is constantly updated to be a good approximation about the current operating point.

Learning happens as long as the gradient estimate, however computed, gives a reasonable descent direction on average — that is, gives direction in which to change the control policy to lower the performance objective value. Operating on inaccurate gradients can result in learning instabilities (distinct from instability in the movement dynamics), as can large gradient steps. This learning instability can be prevented in two different ways. First, when the linear models in equations 5 and 6 are inaccurate, as estimated by their residual being outside of the 95% confidence interval at steady state, the learning rate is set to zero. A second approach to avoiding learning instability is a trust region approach, wherein the maximum gradient-based step-size is limited to a fraction the exploratory noise. We tested both approaches and they give qualitatively similar results.

### Forming motor memories and employing them when useful

We posit a modular memory unit to capture the fact that humans form and maintain memories of previously learned tasks^40^, as opposed to having to re-learn the tasks each time. First, we discuss our model of how such stored ‘motor memories’ are used, and later in this section, we discuss how these motor memories are formed and updated based on experience.

Consider that the human had some past experience in the current task, and used controller parameters *p*_memory_ with associated performance objective values *J*_memory_. We posit that humans move toward this memory with some learning rate as follows:

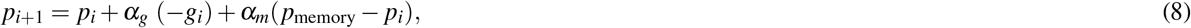

where (*p*_memory_ *− p*_*i*_) is the vector direction toward the memory and *α*_*m*_ is the rate at which memory is approached. We posit that the controller parameters being learned move toward the memory only when *J*_memory_ *< J*_current_, that is, when progressing toward the memory improves the performance. Secondly, to ensure that progress toward memory does not destroy gradient descent even if *J*_memory_ was inaccurately approximated, we posit that the learning rate *α*_*m*_ is modulated via a truncated cosine tuning so that memory is used only when the direction toward memory does not oppose the direction of the negative gradient (Fig. 1d). In *Supplementary Methods*, we elaborate mathematically on why this modulation of the rate toward memory is necessary and sufficient to avoid convergence to a sub-optimal memory.

We conceive of a ‘motor memory’ as a pair of functions *F*_*p*_(*λ*) and *F*_*J*_(*λ*) that output the controller parameters *p*_memory_ = *F*_*p*_(*λ*) and the corresponding performance objective value *J*_memory_ = *F*_*J*_(*λ*) respectively, given the task parameters *λ* (Fig. 1d). The task parameters *λ* could be continuous-valued, for instance, walking speed or assistance level of an exoskeleton, or discrete-valued^41^, for instance, treadmill versus overground walking or presence versus absence of an exoskeleton. As a simple example, the task parameter could be treadmill belt speed *v*_belt_, and the stored motor memory functions *F*_*p*_(*v*_belt_) outputs controller parameters for each walking speed and *F*_*J*_(*v*_belt_) outputs the corresponding performance objective value. In this case, the nervous system could infer the belt speed *v*_belt_ from the sensory stream^55^, by fusing proprioception (which can infer the speed of head relative to foot, *v*_head*/*foot_) and vision (which can sense the speed of head relative to lab, *v*_head*/*lab_), so that the belt speed is given by: *v*_belt_ = *v*_foot*/*lab_ = *v*_head*/*lab_ *−v*_head*/*foot_. For simplicity, we assume that this task parameter inference is independent of any potential perceptual recalibration^22,42^, which is addressed separately in a later section on proprioceptive realignment.

The memory functions *F*_*p*_ and *F*_*J*_ are built over time to approximate the best controllers learned during previous experiences of similar tasks. We posit representations of motor memories via function approximation: in this manuscript, for simplicity, the stored controller parameters *p*_memory_ are linear functions of the task parameters unless otherwise specified, anchored at a nominal tied-belt condition. Such interpolating function approximations for memory are in contrast to discrete memories of experiences without interpolation^2^: these two assumptions have different testable implications for generalization of learning.

The memory functions *F*_*p*_ and *F*_*J*_ have memory parameters *µ*, which determine the function approximation, for instance, the slope and intercept of a linear functions *F*_*P*_(*λ, µ*): here, we allow the slopes of the linear function to change to approximate new experiences, while keeping the intercept at a nominal tied belt walking speed fixed. Analogous to our previous hypothesis that the controller parameters are updated via gradient descent, we posit that these memory parameters are also updated via gradient descent, so that the memory function better approximates controller parameters to be stored. That is, we posit that the nervous system performs: *µ*_i+1_ = *µ*_*i*_*−α*_mf_ ∇_*µ*_ *L*, where *L* is a measure of how well the memory approximates the current controller *p*_*i*_ and *α*_mf_ is the learning rate for memory formation; we used the root mean squared error over all controller parameters being approximated. This memory update happens when the current controller *p*_*i*_ is better than what is already stored in the memory i.e., when *J*_current_ *< J*_memory_ or when the direction toward the memory is not a descent direction. This ensures that memory formation and memory use are mutually exclusive (Fig. 1d).

### Minimal walking biped: dynamics, control, energy, and performance

#### Dynamics

We consider a minimal model of bipedal walking (Supplementary Fig. 1a), consisting of a point-mass upper body and simple legs that can change length and apply forces on the upper body^26,54^. The total metabolic energy cost of walking for this biped is defined as a weighted sum of the positive and negative work done by the stance legs on the upper body and the work done to swing the legs forward^54,95^. For this biped, the periodic energy-optimal walk on solid ground is the inverted pendulum walking gait^26,54,96^, in which the body vaults over the foot in a circular arc on each step (Supplementary Fig. 1b), with the transition from one step to the next achieved via push-off by the trailing leg, followed by a heel-strike at the leading leg (Supplementary Fig. 1c). We use this irreducibly minimal low-dimensional biped model^26,54^ to illustrate the predictive ability of our modeling framework for simplicity and transparency. Further, we show that the simple model is sufficient for explaining the major documented locomotor adaptation phenomena in the literature. The locomotor adaptation modeling framework herein can be generalized to a more complex multibody multimuscle model of a human. The parameters of this biped model were not fit to any data from the adaptation phenomena we seek to explain (see *Supplementary Methods*). When showing metabolic energy cost transients for the model, we show two versions (e.g., in Fig. 2a), one that reflects average metabolic rate over each stride and one that would be measured via indirect calorimetry, which is a low-pass filtered version of the stride-wise cost^97^.

#### Stabilizing feedback control

The biped has two control variables for each leg, namely, step length and push-off magnitude (Supplementary Fig. 1d), for a total of four discrete control actions per stride. These control variables are modulated to keep the biped stable, despite external or internal noisy perturbations and despite a change in the mechanical environment e.g., walking on a split-belt treadmill or with an exoskeleton. The controller keeps the biped stable despite large changes in the body and environment, including external perturbations; this ability to be unaffected by unforeseen changes before any changes to the controller parameters is called robustness, so that the controller is termed ‘robust’^16,17,98^. The values of these control variables on each step are decided by a discrete controller, as described below, derived from our prior human experiments on steady walking^16,17,31^ without fitting any parameters to the data from the adaptation experiments we seek to explain. The body state *s*_*i*_ at midstance at step *i* includes the forward position in the lab frame, the forward velocity in the belt and the lab frame, and the running sum (i.e., discrete integral) of the forward position in the lab frame. The control variables *u*_*i*_ at step *i* are changed by the following linear control rule as a function of the preceding midstance state *s*_*i*_: *u*_*i*_ = *u*_nominal_ + *K* (*s*_*i*_ *−s*_nominal_), where *K* is a matrix of feedback gains^16,31^. The velocity dependence of the control gains ensures that the walker doesn’t fall, the position dependence promotes station-keeping^17,31^, and integral dependence reduces error due to systematic changes in the environment, for instance, changing the treadmill belt speeds or going from a tied to a split treadmill. These terms make the controller a discrete PID controller (proportional-integral-derivative). The nominal periodic motion at each speed is governed by the feedforward push-off and step length values, and these are selected so as to have the same speeds and step lengths as a typical human (Supplementary Fig. 2). The default values for the control gain matrix *K* are then obtained by fitting the dynamics of the model biped to the step to step map of normal human walking on a treadmill^16,17,31,89^. Mathematical details and parameter values are provided in the *Supplementary Methods*. All variables in equations and figures are non-dimensional, unless otherwise mentioned: lengths normalized by leg length *ℓ*, masses normalized by total body mass, and time normalized by 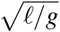, where *g* is acceleration due to gravity^54^.

#### Different locomotor task settings

The biped model described above is expressive enough to capture the different task settings for which we seek to model adaptation: walking with different exoskeleton assistance protocols, at varying belt speeds (tied or split-belt), and with asymmetric leg masses. Here, we briefly describe how the different conditions are simulated by changing the external environment and force it exerts on the biped. See *Supplementary Methods* for mathematical details. The biped model described above allows the individual treadmill belt speeds to be changed as a function of time (Fig. 2a,c). This generality is sufficient to simulate both split-belt and tied-belt treadmill walking conditions. The total metabolic cost computed accounts for individual belt-speed changes because all components of the metabolic cost, namely, push-off work, the heel-strike work, and the leg swing cost are computed by incorporating the relevant belt-speeds and effective leg masses.

For split-belt walking protocols, we usually use non-dimensional belt speeds of 0.5 for the fast belt and 0.25 or 0.3 for the slow belt for many but not all computational results (Figs. 2a, 4, 5, 6,7, 8): these walking speeds and their durations may be different from precise experimental conditions but the qualitative features we illustrate are insensitive to such differences in speeds chosen. We simulated exoskeletons as external devices in parallel to the leg that produce forward forces, or equivalently, ankle torques (Figures 2d and 4e). Walking with periodic exoskeleton input used the perturbation as an additional input state in the controller. For predicting adaptation to foot mass change (Fig. 2b), we incorporated the simplest leg swing dynamics: a point-mass foot, propelled forward with an initial impulse and the foot mass coasting forward passively until heel-strike. For the tied-belt walking, we used the belt speed as the task parameter; for split-belt studies, we used the individual belt speeds as the task parameters; for the added leg mass study, we used the added leg mass as the task parameter. To track gait asymmetry, we use the following two objectives of left-right asymmetry^4^, namely step length asymmetry and step time asymmetry, defined as follows:

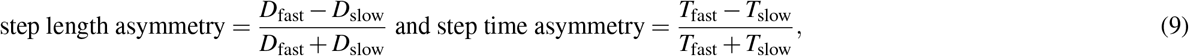

where the fast and the slow step lengths (*D*_fast_ and *D*_slow_) are defined at heel strike as in Supplementary Fig. 1e and the step times *T*_fast_ and *T*_slow_ are the stance times when on the fast and slow belts respectively. We use analogous objectives when the biped walks with an asymmetric foot mass. There can be other ways of quantifying asymmetry and we chose asymmetry objectives that are commonly used to empirically track adaptation^4,5,45^. A zero value indicates symmetry with respect to these measures, but does not imply perfect left-right symmetry of the entire motion.

All computational work was performed in MATLAB (version 2022a). See Supplementary Methods and our codebase implementing these simulations, *LocAd*^30^, shared via a public repository. Data from prior manuscripts^5,6,8,18,20,28,45,48,51,99,100^ are plotted in Figures 2, 6a, and 7b and Supplementary Figures 2, 4, 6, as cited in place, to illustrate how model-based predictions agree qualitatively with experimental results in prior studies.

### Alternative to performance optimization: Comparison with proprioceptive realignment

Experimental evidence during split-belt adaptation suggests some recalibration of proprioception by the two legs^42^ and has been argued to be at least partially responsible for kinematic adaptation^22,42^ based on correlation of timescales between such realignment and adaptation. No mathematical model been been previously proposed for how such realignment may happen. Without such a mathematical model, it is impossible to know whether the direction of adaptation due to such realignment will be consistent with or opposing that observed in experiment. Here, we first present such a mathematical model and then test the extent to which it explains adaptation on a split-belt treadmill. We implement this proprioceptive realignment as a sensory recalibration of the input to the stabilizing controller, replacing the gradient-based reinforcement learner (Fig. 9a-b).

Recalibration takes place when there is substantial conflict between what is expected by the nervous system and what is sensed^11^. The key missing hypothesis in extending such sensory recalibration to locomotor adaptation lies in the question: what error is the nervous system using to drive recalibration during locomotion? We hypothesize that, given the typical walking experienced in daily life, the nervous system expects the two legs to be on a common surface: this expectation results in a sensory conflict on a split-belt treadmill with both feet experiencing unequal belt speeds.

When the walking surface has fixed speed and the visual environment is uniform, the walking speed can be estimated by the nervous system by two sensory modalities^101^: vision (based on visual flow) and proprioception (by integrating joint angles and angle rates from muscle spindles and Golgi tendon organs). On a treadmill in a lab, vision has information about how the head moves with respect to the lab, so we identify the visual speed with *v*_body*/*lab_. Proprioception has information about how fast the body parts move relative to the stance foot on the belt, so we identify proprioception with *v*_body*/*belt_. Thus, the body has information to implicitly estimate the belt speed via the following equation: *v*_belt*/*lab_ = *v*_body*/*lab_ *− v*_body*/*belt_. On a split-belt treadmill, all these speeds will be belt-specific, e.g., *v*_belt,1*/*lab_ and *v*_belt,2*/*lab_. The expectation that both legs contact a common surface can be expressed as the equality of these individual belt speeds: *v*_belt,1*/*lab_ = *v*_belt,2*/*lab_. We posit that deviations from this equality result in slow recalibration. Consistent with much of the arm reaching literature, we recalibrate only the proprioceptive sense, hence the term proprioceptive realignment.

Say, 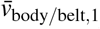 and, 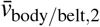 are the proprioceptively obtained sensory information from the two legs without recalibration, and 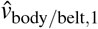 and 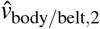 are the recalibrated versions. The two versions are related by:

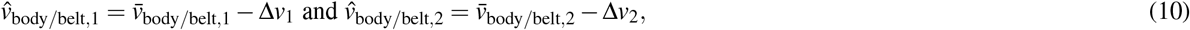

where Δ*v*_1_ and Δ*v*_2_ are the recalibrative corrections. We describe this recalibration as happening via a state observer with two timescales: a fast timescale process estimating a common belt speed 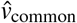 using proprioceptive information from both legs and a slow timescale process estimating the recalibrative corrections Δ*v*_1_ and Δ*v*_2_ for each leg separately. The common belt speed estimate is updated every step *j* via a state observer as follows:

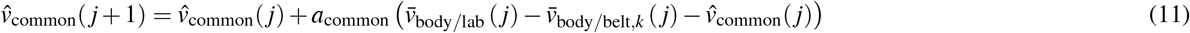

where *k* equals 1 or 2 for odd and even step number given by *j*, respectively, and *a*_common_ is a rate constant proportional to the time spent on each step – but we treat as constant for simplicity. This equation results in convergence of 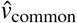 to the average belt speed. The recalibration Δ*v*_*k*_ is the current estimated perturbation of the individual belt speed from the estimated common speed 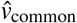 This recalibration is updated on every stride *i* as: 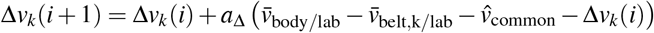 Here, the rate constant *a*_Δ_ is much smaller than *a*_common_, reflecting the slower timescale at which the perturbation estimate 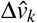 is updated. Here, 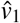 is updated on odd steps and 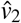 is updated on even steps. This perturbation estimate Δ*v*_*k*_ eventually converges to the deviation of the actual belt speed from the common speed. These state observer equations for recalibration are similar to estimating the belt speed via a state estimator reflecting an expectation that tied-belt changes are much more likely than split-belt changes, modeled by the noise covariance matrix for belt speed changes having large diagonal elements (governing co-variation of belt speeds) and small off-diagonal elements (governing belt speed differences). Further, while we have introduced the latent variables *v*_common_ and Δ*v*_*k*_ in the above description, the recalibration equations can be written without such latent variables. The recalibrated proprioceptive information 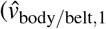 and 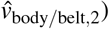 is used in the stabilizing controller instead of the direct proprioception. In the Results, we show effects of 50% and 100% recalibration: 100% corresponds to using the full correction Δ*v*_*k*_ and 50% uses 0.5 Δ*v*_*k*_ in the recalibration equation 10.

### Alternative to energy minimization: Comparison with reducing kinematic task error

Minimizing kinematic task error first requires defining what the desired or expected kinematics are. To define this, we first note that slow timescale error minimization is not thought to underlie tied-belt walking adaptation, or at least the timescales of adaptation to tied-belt speed changes are much faster than for split-belt adaptation^28^. So, we posit that the nervous system treats tied-belt walking — or walking on the same surface with both legs — as the normal state of affairs, basing the desired kinematics on an implicit assumption of tied-belt walking. The nervous system estimates the common belt speed 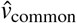 as in the previous section, using visual flow and proprioception from both legs (see eq. 11 and Fig. 8b). This common belt speed 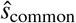 is then used to make a prediction for the body midstance state ŝ_common_ based on memory, which is compared with actual sensory information 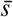 to compute the task error 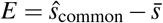.

In simple tasks such as arm reaching, where both the error and the action is one-dimensional, it is possible to reduce error via a simple ‘error-based learning model’ with a single or dual rate process^46^ or via learning rate adaptation^9^. However, in tasks such as walking in a novel environment, because body state and the number of actuators is high-dimensional, the nervous system may not have a priori inverse model to produce the motor actions that reduce the error. So, a simple one-dimensional error based learning model is not appropriate. We instead model the error reduction as proceeding analogous to energy reduction via gradient descent using exploratory variability to estimate gradients, as a special case of the framework in Fig. 1. The kinematic task error being minimized may also be considered a kind of sensory prediction error, as the error from the kinematic state predicted or expected by the nervous system given the belt speed estimate.

### Interaction with explicit control

The mechanisms proposed herein are for implicit adaptation, but these mechanisms allow for explicit (conscious) control acting in parallel to implicit adaptation. We show how our model can be extended to interact with explicit input by implementing an visually-informed explicit control module for the reduction of step length asymmetry^10^. On each step, the explicit control module outputs a correction to the desired step length proportional to the step length asymmetry on the previous stride, with the intention of reducing the step length asymmetry on the current step. This output from the explicit control module is added to the desired step length output from the implicit adaptation module (the default adaptation mechanisms here), so that the net total control input is used by the stabilizing controller. The additive and parallel nature of the implicit and explicit modules are as proposed for explicit control in arm reaching studies^23^. The architecture of the interaction between the implicit adaptation and explicit control is such that the implicit module is only aware of its own output and not that of the explicit module; thus, the implicit module optimizes the objective with respect to its own output.

### Prospective experiments

The computational model we put forth here can be used to design prospective experiments, augmenting experimenter intuition. Here, we conducted two model-guided experiments to test predictions of the model that are surprising when compared to the existing literature: (1) on the effect of environment noise on locomotor adaptation, and (2) on the effect of an immediately preceding counter-perturbation on a subsequent adaptation.

Twenty five participants (19 male, 6 female, self-reported sex, age 21.9 *±* 3 years, mean *±* s.d.) participated with informed consent and the experiments were approved by the Ohio State University IRB. Participants were assigned randomly into two groups: sixteen participants performed experiment 1 (12 male, 4 female, age 21.7 *±* 3 years) and nine participants (6 male, 3 female, age 22.3 *±* 3 years) performed experiment 2. Both experiments involved walking on a split-belt treadmill (Bertec Inc.), with the details of the protocol provided below. Foot movement was tracked via a Vicon T20 motion capture system (Vicon Nexus 1.x). Sex or age were not used as an explanatory variable in any analysis, as the computational model tested does not include such variables.

Experiment 1 was designed to test the model prediction that when the belt noise level was sufficiently high, learning can be degraded, which is surprising relative to a prior finding that a modest level of belt noise can slightly enhance learning as measured by after-effects^20^. For this experiment, the participants were sub-divided into two groups of eight: one group performed a no-noise abrupt protocol (Fig. 7a), in which participants started walking under tied-belt conditions at 0.9 m/s, then adapted to split-belt condition of 0.6 m/s and 1.2 m/s kept constant for 10 minutes, followed by three minutes of tied-belt walking at 0.6 m/s; the second group had an identical protocol except the split-belt condition involved continuously changing belt speed for just the fast belt, fluctuating in a piecewise linear manner with zero mean and 0.2 m/s standard deviation (normally distributed). The consecutive grid points of the piecewise linear noise were separated by 1.2 seconds, roughly equal to a stride period, so that the noise value was different two strides apart (the noise in^20^ was changed every 3 seconds, and thus had greater temporal correlation); speed changes had 0.1-0.2 m/s^2^ accelerations. The noise standard deviation was set at a lower level in simulation (0.04 m/s) to ensure stability. The post-adaptation after-effect in step length asymmetry after baseline subtraction, averaged over the first 8 strides (about 10 seconds) was used as a measure of adaptation similar to prior work^20^. We compared these after-effects between the noise and no-noise case, testing the hypothesis that the noise case has lower after-effects.

Experiment 2 was designed to test the model prediction regarding savings, specifically whether experiencing a counter perturbation B beforehand, interferes with adaptation to perturbation A. Previous experiment had found that if B and A were separated by a washout period, the adaptation to A was not significantly affected compared to not having experienced B. Our model had a distinct prediction for when A immediately followed B, without a washout period W. So, participants performed this experimental protocol T-B-A (Fig. 6) in which 4 minutes of walking on a tied-belt at 0.9 m/s (T) was followed by a split-belt condition with belt speeds of 0.6 and 1.2 m/s for 10 minutes (B), immediately followed by the opposite split-belt condition 1.2 and 0.6 m/s for 10 minutes. Equivalent to comparing the A of protocols T-A and T-B-A by symmetry, we compared the initial transient and the time-constant of the two adaptation periods B and A in T-B-A: the B of T-B-A was without prior split-belt experience and the A of T-B-A is the adaptation phase just after a counter-perturbation.

## Supporting information

Supplementary Appendic

## Data availability

The experimental data generated in this study have been deposited in the Dryad database with DOI: 10.5061/dryad.kh18932gq. Source data are provided with this paper. Other human experimental data or results referred to herein are available in previously published manuscripts^5,6,8,18,20,28,45,48,51,52,99,100^.

## Code availability

Code associated with this paper, *LocAd*^30^, is available at: https://github.com/SeethapathiLab/LocAd

## Acknowledgements

N.S. was funded by the Massachusetts Institute of Technology. M.S. was supported in part by NIH grant R01GM135923-01 and NSF SCH grant 2014506.

## Author contributions statement

NS initially conceived the central question. NS and MS conceived the theoretical framework, created mathematical models, performed computations, analyzed the results, and wrote the manuscript. BC contributed additional models and calculations. All authors reviewed and approved the manuscript.

## Additional information

**Supplementary Methods** is available for this paper.

## References

1. Day, K. A., Leech, K. A., Roemmich, R. T. & Bastian, A. J. Accelerating locomotor savings in learning: compressing four training days to one. J. Neurophys. 119, 2100–2113 (2018).

2. Selinger, J. C., Wong, J. D., Simha, S. N. & Donelan, J. M. How humans initiate energy optimization and converge on their optimal gaits. J. Exp. Biol. 222, jeb198234 (2019).

3. Bastian, A. J. Learning to predict the future: the cerebellum adapts feedforward movement control. Curr. Opin. Neurobiol. 16, 645–649 (2006).

4. Reisman, D. S., Block, H. J. & Bastian, A. J. Interlimb coordination during locomotion: what can be adapted and stored? J. Neurophysiol. 94, 2403–2415 (2005).

5. Noble, J. W. & Prentice, S. D. Adaptation to unilateral change in lower limb mechanical properties during human walking. Exp. Brain Res. 169, 482–495 (2006).

6. Finley, J., Bastian, A. & Gottschall, J. Learning to be economical: the energy cost of walking tracks motor adaptation. J. Physiol. 591, 1081–1095 (2013).

7. Sánchez, N., Simha, S. N., Donelan, J. M. & Finley, J. M. Taking advantage of external mechanical work to reduce metabolic cost: The mechanics and energetics of split-belt treadmill walking. J. Physiol. 597, 4053–4068 (2019).

8. Selinger, J. C. O, Connor, S. M., Wong, J. D. & Donelan, J. M. Humans can continuously optimize energetic cost during walking. Curr. Biol. 25, 2452–2456 (2015).

9. Herzfeld, D. J., Vaswani, P. A., Marko, M. K. & Shadmehr, R. A memory of errors in sensorimotor learning. Science 345, 1349–1353 (2014).

10. Roemmich, R. T., Long, A. W. & Bastian, A. J. Seeing the errors you feel enhances locomotor performance but not learning. Curr. biology 26, 2707–2716 (2016).

11. Izawa, J. & Shadmehr, R. Learning from sensory and reward prediction errors during motor adaptation. PLoS computational biology 7, e1002012 (2011).

12. Tseng, Y.-w., Diedrichsen, J., Krakauer, J. W., Shadmehr, R. & Bastian, A. J. Sensory prediction errors drive cerebellum-dependent adaptation of reaching. J. neurophysiology 98, 54–62 (2007).

13. Therrien, A. S., Wolpert, D. M. & Bastian, A. J. Effective reinforcement learning following cerebellar damage requires a balance between exploration and motor noise. Brain 139, 101–114 (2016).

14. Hof, A., Vermerris, S. & Gjaltema, W. Balance responses to lateral perturbations in human treadmill walking. J Exp Biol 213, 2655–2664 (2010).

15. Seyfarth, A., Geyer, H. & Herr, H. Swing-leg retraction: a simple control model for stable running. J Exp Biol 206, 2547–2555 (2003).

16. Joshi, V. & Srinivasan, M. A controller for walking derived from how humans recover from perturbations. J. Roy. Soc. Interface 16, 20190027 (2019).

17. Seethapathi, N. & Srinivasan, M. Step-to-step variations in human running reveal how humans run without falling. Elife 8, e38371 (2019).

18. Roemmich, R. T. & Bastian, A. J. Two ways to save a newly learned motor pattern. J. Neurophys. 113, 3519–3530 (2015).

19. Simha, S. N., Wong, J. D., Selinger, J. C., Abram, S. J. & Donelan, J. M. Increasing the gradient of energetic cost does not initiate adaptation in human walking. J. Neurophysiol. 126, 440–450 (2021).

20. Torres-Oviedo, G. & Bastian, A. J. Natural error patterns enable transfer of motor learning to novel contexts. J. Neurophysiol. 107, 346–356 (2012).

21. Long, A. W., Roemmich, R. T. & Bastian, A. J. Blocking trial-by-trial error correction does not interfere with motor learning in human walking. J. neurophysiology 115, 2341–2348 (2016).

22. Rossi, C., Bastian, A. J. & Therrien, A. S. Mechanisms of proprioceptive realignment in human motor learning. Curr. Opin. Physiol. 20, 186–197 (2021).

23. Taylor, J. A. & Ivry, R. B. Flexible cognitive strategies during motor learning. PLoS computational biology 7, e1001096 (2011).

24. Fujiki, S. et al. Adaptation mechanism of interlimb coordination in human split-belt treadmill walking through learning of foot contact timing: a robotics study. J. Royal Soc. Interface 12, 20150542 (2015).

25. Seethapathi, N. Transients, Variability, Stability and Energy in Human Locomotion. Ph.D. thesis, The Ohio State University (2018).

26. Srinivasan, M. & Ruina, A. Computer optimization of a minimal biped model discovers walking and running. Nature 439, 72–75 (2006).

27. Simha, S. N., Wong, J. D., Selinger, J. C., Abram, S. J. & Donelan, J. M. Increasing the gradient of energetic cost does not initiate adaptation in human walking. bioRxiv (2020).

28. Pagliara, R., Snaterse, M. & Donelan, J. M. Fast and slow processes underlie the selection of both step frequency and walking speed. J. Exp. Biol. 217, 2939–2946 (2014).

29. Ahn, J. & Hogan, N. A simple state-determined model reproduces entrainment and phase-locking of human walking. PloS one 7, e47963 (2012).

30. Seethapathi, N., Clark, B. & Srinivasan, M. Locad: Code for ‘exploration-based learning of a stabilizing controller predicts locomotor adaptation’. https://github.com/SeethapathiLab/LocAd (2024).

31. Wang, Y. & Srinivasan, M. Stepping in the direction of the fall: the next foot placement can be predicted from current upper body state in steady-state walking. Biol. Lett. 10, 20140405 (2014).

32. Ralston, H. J. Energy-speed relation and optimal speed during level walking. Int Z angew Physiol einschl Arbeitsphysiol 17, 277–283 (1958).

33. Zarrugh, M., Todd, F. & Ralston, H. Optimization of energy expenditure during level walking. Eur. J. Appl. Physiol. Occup. Physiol. 33, 293–306 (1974).

34. Long, L. L. & Srinivasan, M. Walking, running, and resting under time, distance, and average speed constraints: optimality of walk–run–rest mixtures. J. R. Soc. Interface 10, 20120980 (2013).

35. Seethapathi, N. & Srinivasan, M. The metabolic cost of changing walking speeds is significant, implies lower optimal speeds for shorter distances, and increases daily energy estimates. Biol. Lett. 11, 20150486 (2015).

36. Bertram, J. & Ruina, A. Multiple walking speed–frequency relations are predicted by constrained optimization. J. Theor. Biol. 209, 445–453 (2001).

37. Torres-Oviedo, G. & Bastian, A. J. Seeing is believing: effects of visual contextual cues on learning and transfer of locomotor adaptation. J. Neurosci. 30, 17015–17022 (2010).

38. Wu, H. G., Miyamoto, Y. R., Castro, L. N. G., Ölveczky, B. P. & Smith, M. A. Temporal structure of motor variability is dynamically regulated and predicts motor learning ability. Nat. neuroscience 17, 312–321 (2014).

39. Sutton, R. S. & Barto, A. G. Reinforcement learning: An introduction (MIT press, 2018).

40. Brashers-Krug, T., Shadmehr, R. & Bizzi, E. Consolidation in human motor memory. Nature 382, 252–255 (1996).

41. Heald, J. B., Lengyel, M. & Wolpert, D. M. Contextual inference underlies the learning of sensorimotor repertoires. Nature 600, 489–493 (2021).

42. Vazquez, A., Statton, M. A., Busgang, S. A. & Bastian, A. J. Split-belt walking adaptation recalibrates sensorimotor estimates of leg speed but not position or force. J. neurophysiology 114, 3255–3267 (2015).

43. Kluzik, J., Diedrichsen, J., Shadmehr, R. & Bastian, A. J. Reach adaptation: what determines whether we learn an internal model of the tool or adapt the model of our arm? J. neurophysiology 100, 1455–1464 (2008).

44. Choi, J. T., Vining, E. P., Reisman, D. S. & Bastian, A. J. Walking flexibility after hemispherectomy: split-belt treadmill adaptation and feedback control. Brain 132, 722–733 (2008).

45. Sánchez, N., Simha, S. N., Donelan, J. M. & Finley, J. M. Using asymmetry to your advantage: learning to acquire and accept external assistance during prolonged split-belt walking. J. Neurophysiol. 125, 344–357 (2021).

46. Smith, M. A., Ghazizadeh, A. & Shadmehr, R. Interacting adaptive processes with different timescales underlie short-term motor learning. PLoS Biol 4, e179 (2006).

47. Stenum, J. & Choi, J. T. Step time asymmetry but not step length asymmetry is adapted to optimize energy cost of split-belt treadmill walking. The J. Physiol. 598, 4063–4078 (2020).

48. Ochoa, J., Sternad, D. & Hogan, N. Treadmill vs. overground walking: different response to physical interaction. J. Neurophysiol. 118, 2089–2102 (2017).

49. Buurke, T. J., Lamoth, C. J., van der Woude, L. H. & den Otter, R. Handrail holding during treadmill walking reduces locomotor learning in able-bodied persons. IEEE Transactions on Neural Syst. Rehabil. Eng. 27, 1753–1759 (2019).

50. Park, S. & Finley, J. M. Manual stabilization reveals a transient role for balance control during locomotor adaptation. J. Neurophysiol. 128, 808–818 (2022).

51. Malone, L. A., Vasudevan, E. V. & Bastian, A. J. Motor adaptation training for faster relearning. J. Neurosci. 31, 15136–15143 (2011).

52. Leech, K. A., Roemmich, R. T. & Bastian, A. J. Creating flexible motor memories in human walking. Sci. Rep. 8, 1–10 (2018).

53. Lam, J. et al. Impaired implicit learning and feedback processing after stroke. Neurosci. 314, 116–124 (2016).

54. Srinivasan, M. Fifteen observations on the structure of energy-minimizing gaits in many simple biped models. J. R. Soc. Interface 8, 74–98 (2011).

55. Srinivasan, M. Optimal speeds for walking and running, and walking on a moving walkway. Chaos 19, 026112 (2009).

56. Handford, M. L. & Srinivasan, M. Robotic lower limb prosthesis design through simultaneous computer optimizations of human and prosthesis costs. Sci. Rep. 6, 19983 (2016).

57. Henriques, D. Y. & Cressman, E. K. Visuomotor adaptation and proprioceptive recalibration. J. motor behavior 44, 435–444 (2012).

58. Tsay, J. S., Kim, H., Haith, A. M. & Ivry, R. B. Understanding implicit sensorimotor adaptation as a process of proprioceptive re-alignment. Elife 11, e76639 (2022).

59. Reisman, D., Wityk, R., Silver, K. & Bastian, A. Locomotor adaptation on a split-belt treadmill can improve walking symmetry post-stroke. Brain 130, 1861–1872 (2007).

60. Leech, K. A., Day, K. A., Roemmich, R. T. & Bastian, A. J. Movement and perception recalibrate differently across multiple days of locomotor learning. J. neurophysiology 120, 2130–2137 (2018).

61. Friston, K. The free-energy principle: a unified brain theory? Nat. reviews neuroscience 11, 127–138 (2010).

62. Abram, S. J., Selinger, J. C. & Donelan, J. M. Energy optimization is a major objective in the real-time control of step width in human walking. J. biomechanics 91, 85–91 (2019).

63. Wong, J. D., Selinger, J. C. & Donelan, J. M. Is natural variability in gait sufficient to initiate spontaneous energy optimization in human walking? J. neurophysiology 121, 1848–1855 (2019).

64. Pekny, S. E., Criscimagna-Hemminger, S. E. & Shadmehr, R. Protection and expression of human motor memories. J. Neurosci. 31, 13829–13839 (2011).

65. Zarahn, E., Weston, G. D., Liang, J., Mazzoni, P. & Krakauer, J. W. Explaining savings for visuomotor adaptation: linear time-invariant state-space models are not sufficient. J. neurophysiology 100, 2537–2548 (2008).

66. Witten, E. String theory dynamics in various dimensions. Nucl. Phys. B 443, 85–126 (1995).

67. Anderson, B. D. Failures of adaptive control theory and their resolution. Comm. Info. Sys. 5, 1–20 (2005).

68. Hamilton, A. & Grafton, S. T. The motor hierarchy: from kinematics to goals and intentions. Sensorimotor foundations higher cognition 22, 381–408 (2007).

69. Armstrong, D. Supraspinal contributions to the initiation and control of locomotion in the cat. Prog. neurobiology 26, 273–361 (1986).

70. Drew, T., Prentice, S. & Schepens, B. Cortical and brainstem control of locomotion. Prog. brain research 143, 251–261 (2004).

71. Statton, M. A., Vazquez, A., Morton, S. M., Vasudevan, E. V. & Bastian, A. J. Making sense of cerebellar contributions to perceptual and motor adaptation. The Cerebellum 17, 111–121 (2018).

72. Morton, S. M. & Bastian, A. J. Cerebellar contributions to locomotor adaptations during splitbelt treadmill walking. J. Neurosci. 26, 9107–9116 (2006).

73. Bastian, A. J. Moving, sensing and learning with cerebellar damage. Curr. opinion neurobiology 21, 596–601 (2011).

74. Darmohray, D. M., Jacobs, J. R., Marques, H. G. & Carey, M. R. Spatial and temporal locomotor learning in mouse cerebellum. Neuron 102, 217–231 (2019).

75. Marr, D. Vision: A computational investigation into the human representation and processing of visual information (MIT press, 2010).

76. Therrien, A. S., Statton, M. A. & Bastian, A. J. Reinforcement signaling can be used to reduce elements of cerebellar reaching ataxia. The Cerebellum 20, 62–73 (2021).

77. Todorov, E. & Jordan, M. I. Optimal feedback control as a theory of motor coordination. Nat. Neurosci. 5, 1226–1235 (2002).

78. Jordan, M. I. & Rumelhart, D. E. Forward models: Supervised learning with a distal teacher. Cog. Sci. 16, 307–354 (1992).

79. Peng, X. B., Berseth, G., Yin, K. & Van De Panne, M. Deeploco: Dynamic locomotion skills using hierarchical deep reinforcement learning. ACM Trans. Graph. (TOG) 36, 1–13 (2017).

80. Kidziński, Ł. et al. Learning to run challenge solutions: Adapting reinforcement learning methods for neuromusculoskeletal environments. In The NIPS’17 Competition: Building Intelligent Systems, 121–153 (Springer, 2018).

81. Xie, Z., Berseth, G., Clary, P., Hurst, J. & van de Panne, M. Feedback control for cassie with deep reinforcement learning. In 2018 IEEE/RSJ International Conference on Intelligent Robots and Systems (IROS), 1241–1246 (IEEE, 2018).

82. Ajemian, R. & Hogan, N. Experimenting with theoretical motor neuroscience. J. motor behavior 42, 333–342 (2010).

83. Cashaback, J. G., McGregor, H. R., Mohatarem, A. & Gribble, P. L. Dissociating error-based and reinforcement-based loss functions during sensorimotor learning. PLoS computational biology 13, e1005623 (2017).

84. Albert, S. T. et al. Competition between parallel sensorimotor learning systems. Elife 11, e65361 (2022).

85. Sombric, C. J., Calvert, J. S. & Torres-Oviedo, G. Large propulsion demands increase locomotor adaptation at the expense of step length symmetry. Front. physiology 10, 60 (2019).

86. Zhang, J. et al. Human-in-the-loop optimization of exoskeleton assistance during walking. Science 356, 1280–1284 (2017).

87. Vasudevan, E. V. & Bastian, A. J. Split-belt treadmill adaptation shows different functional networks for fast and slow human walking. J. neurophysiology 103, 183–191 (2010).

88. Gordon, K. E. & Ferris, D. P. Learning to walk with a robotic ankle exoskeleton. J. Biomech. 40, 2636–2644 (2007).

89. Perry, J. A. & Srinivasan, M. Walking with wider steps changes foot placement control, increases kinematic variability and does not improve linear stability. Roy. Soc. Open Sci. 4, 160627 (2017).

90. Goodwin, G. C. & Sin, K. S. Adaptive filtering prediction and control (Courier Corporation, 2014).

91. Herzfeld, D. J. & Shadmehr, R. Motor variability is not noise, but grist for the learning mill. Nat. Neurosci. 17, 149–150 (2014).

92. Harris, C. M. & Wolpert, D. M. Signal-dependent noise determines motor planning. Nature 394, 780–784 (1998).

93. Osborne, L. C., Lisberger, S. G. & Bialek, W. A sensory source for motor variation. Nature 437, 412–416 (2005).

94. Sutton, R. S., McAllester, D. A., Singh, S. P. & Mansour, Y. Policy gradient methods for reinforcement learning with function approximation. In Adv. Neur. Info. Proc. Sys., 1057–1063 (2000).

95. Kuo, A. A simple model of bipedal walking predicts the preferred speed–step length relationship. J. Biomech. Eng. 123, 264–269 (2001).

96. Srinivasan, M. & Ruina, A. Idealized walking and running gaits minimize work. Proc. Roy. Soc. A 463, 2429–2446 (2007).

97. Selinger, J. C. & Donelan, J. M. Estimating instantaneous energetic cost during non-steady-state gait. J. Appl. Physiol. 117, 1406–1415 (2014).

98. Zhou, K. & Doyle, J. C. Essentials of robust control, vol. 104 (Prentice hall Upper Saddle River, NJ, 1998).

99. Bertram, J. & Ruina, A. Multiple walking speed-frequency relations are predicted by constrained optimization. J. theor. Biol. 209, 445–453 (2001).

100. Minetti, A. & Alexander, R. A theory of metabolic costs for bipedal gaits. J. Theor. Biol. 186, 467–476 (1997).

101. Srinivasan, M. Optimal speeds for walking and running, and walking on a moving walkway. CHAOS 19, 026112 (2009).

