## Supplementary Appendic for "Exploration-based learning of a stabilizing controller predicts locomotor adaptation"

### Supplementary Information for “Exploration-based learning of a stabilizing controller predicts locomotor adaptation”

#### Supplementary Figures

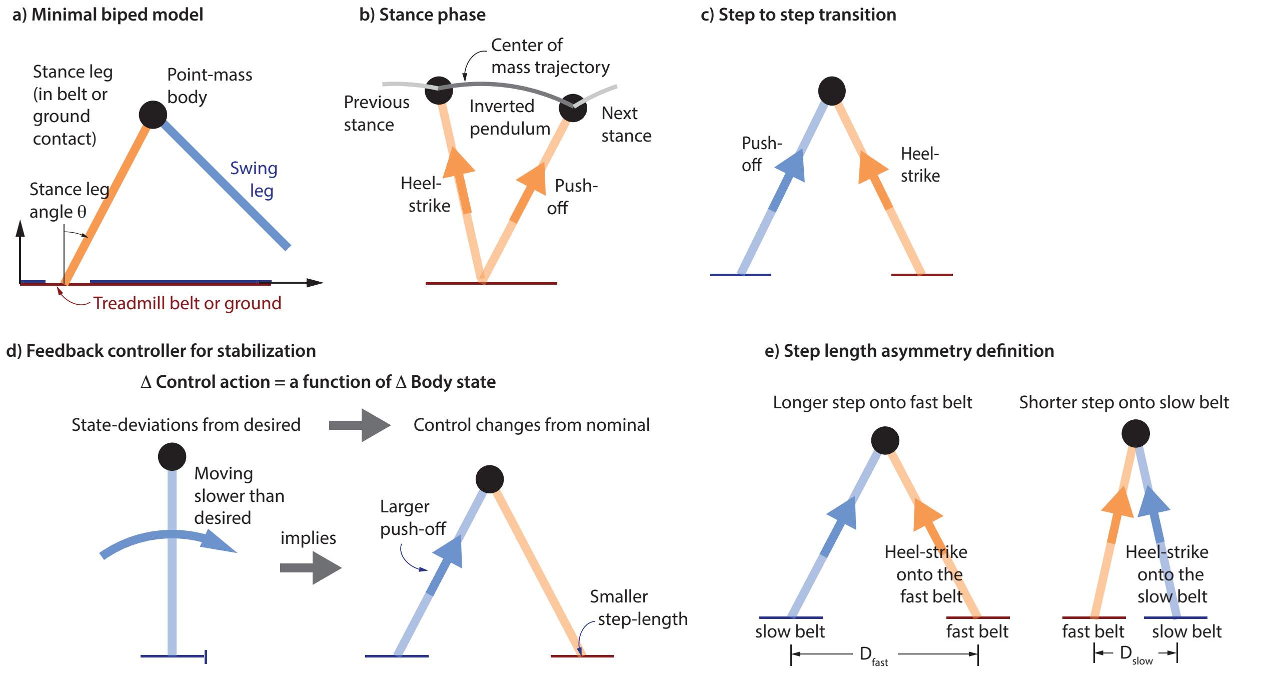


**Supplementary Figure 1. Biped model and controller. a)** A minimal biped model with a point-mass upper body, **b)** using an inverted pendulum walking gait, **c)** with step to step transitions mediated by impulsive push-offs and heel-strikes[1,2]. **d)** The push-off and step lengths on every step are controlled via state feedback on center of mass state, so that moving faster than desired or being behind the desired position results in a larger push-off and a smaller step length[3–5]. **e)** The step length asymmetry is computed based on step lengths $D_{\text{fast}}$ and $D_{\text{slow}}$ (distance between the feet) measured at the time of heel-strike onto the fast or slow belt in the case of split-belt walking. For walking with an asymmetric foot mass, an analogous step length asymmetry is computed by using step lengths $D_{\text{heavy}}$ and $D_{\text{normal}}$ onto the heavier foot and the normal foot respectively. Source data are provided as a Source Data file.

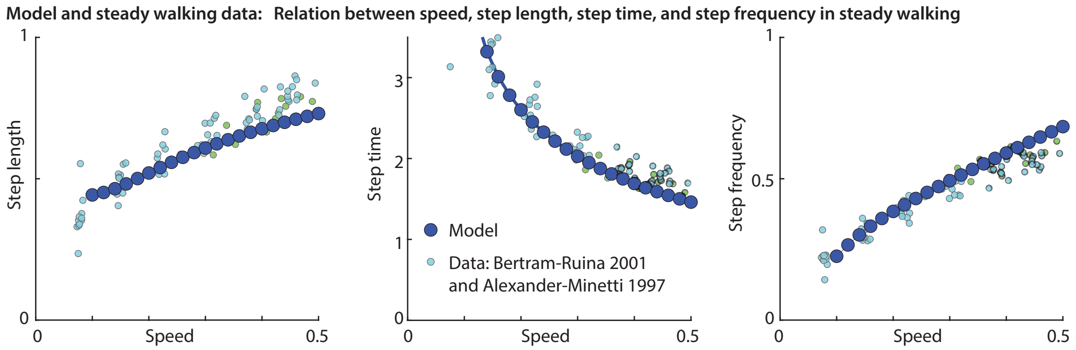


**Supplementary Figure 2. Steady state walking characteristics.** At each steady walking speed, humans use a stereotypical step length — or equivalently, step frequency and step time, derivable from speed and step length[6,7]. One parameter of the biped model (scaling of swing cost) is chosen such that the optimal gait captures how the steady step length, step frequency, and step time change with speed in human data[6,7].

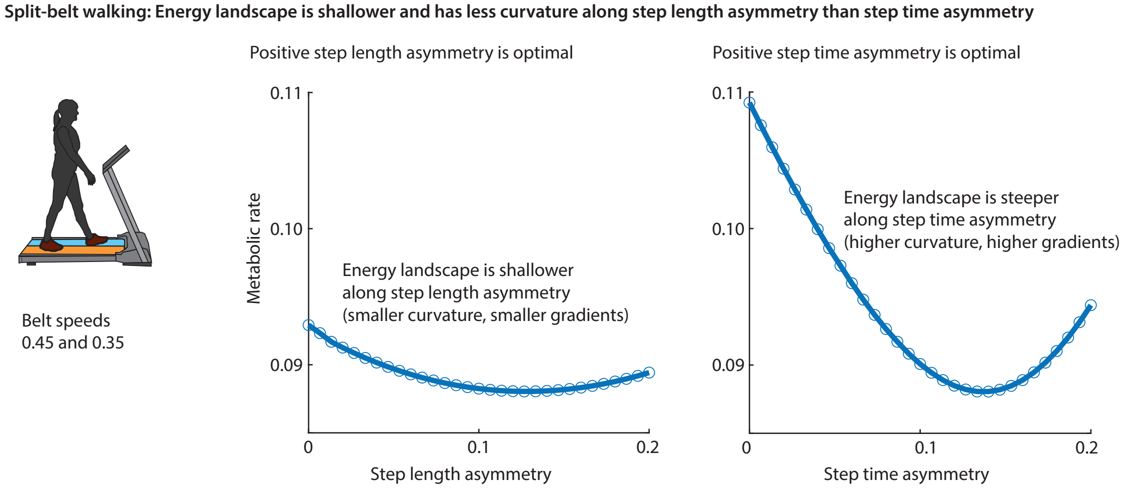


**Supplementary Figure 3. Step length vs step time asymmetries.** For walking on a split-belt treadmill, the energy landscape has optimum at positive step length asymmetry and positive step time asymmetry. The curvature and thus the slopes are higher along the step time asymmetry than the step length asymmetry directions, thus predicting that gradient descent-based learning will converge faster in step time asymmetry rather than step length asymmetry. Further, the relative flatness of the optimum along step length asymmetry compared to along step time asymmetry means the behavior can be further from the optimum along step length asymmetry direction for a given energy penalty. So, if the learning does not converge fully, we expect it to be further away from the optimum along the step length asymmetry compared to step time asymmetry. Finally, note that a given step length or step time asymmetry can be chosen in infinitely many ways, for instance, with different total or average step lengths; so this would need to be a higher dimensional plot to get a better characterization of the energy landscape. We have shown two slices of this higher dimensional cost landscape to compare the curvatures along two directions in the units chosen; the slices are chosen through the optimum, with the other free variables chosen to be their optimal value given the appropriate step length or step time asymmetry value. Source data are provided as a Source Data file.

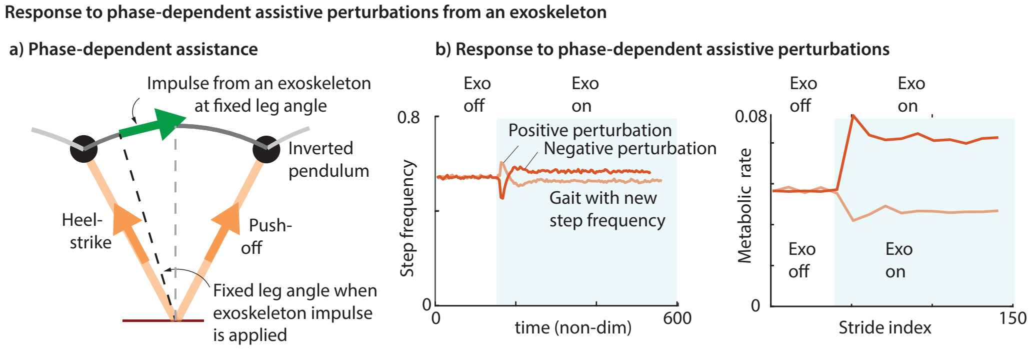


**Supplementary Figure 4.** **State-dependent exoskeleton assistance. a)** Forward assistive torques are applied by the exoskeleton starting at fixed leg angles, with the torque being angle dependent. This makes the assistance periodic in gait phase defined based on leg angle. **b)** The reinforcement learner learns to take advantage of the assistive pulls by changing the step timing slightly, with the transient being due to both the stabilizing feedback controller and the reinforcement learner. Period when the exoskeleton is providing perturbations is shaded light blue. Perturbations were applied at leg angle equalling -0.1 radians, where an angle of 0 corresponds to a vertical leg. Accelerating and decelerating perturbations both result in periodic gaits, but with periods lower or higher than the nominal gait period. Source data are provided as a Source Data file.

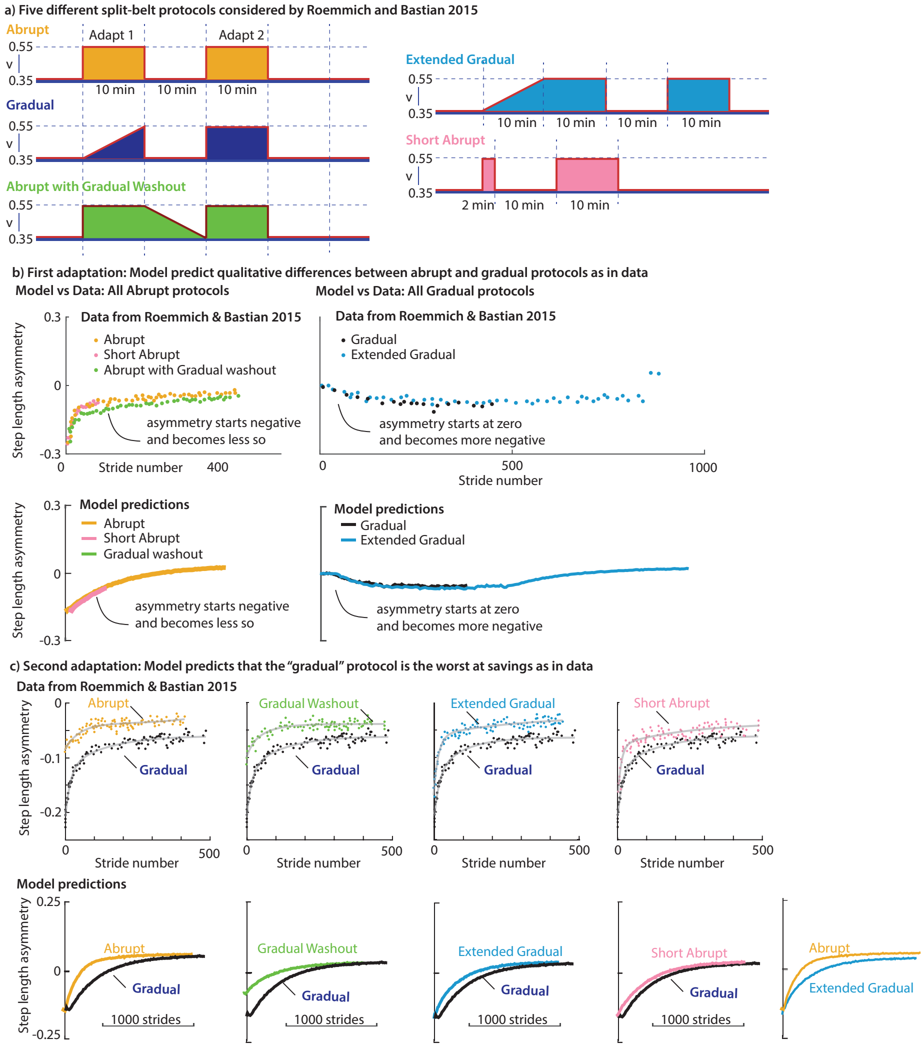


**Supplementary Figure 5. Protocols with abrupt, gradual, and extended adaptation regimes. a)** We simulated the different adaptation protocols consisting of gradual and abrupt introduction of a split-belt condition, different extents of the adaptation regime, and gradual versus abrupt ending to the adaptation regime, analogous to those considered by Roemmich and Bastian[8]. Different speed change protocols have different color shadings, and the behavior in panels b and c use the same colors. **b)** During first adaptation, the protocols that begin with abrupt change in belt speeds showed qualitatively different transients compared to the protocols with gradual changes in belt speeds: specifically, the abrupt adaptation phases had an initial negative step length asymmetry that slowly became less negative; the gradual adaptation phases had an initial zero step length asymmetry that slowly became more negative. These qualitative differences in experimental data were predicted by the model for appropriate model parameters. **c)** The model predicts that the second adaptation of every other protocol will have a smaller initial transient and more savings than 'gradual' protocol, as is qualitatively seen in the experimental data. Further, the longer duration abrupt first adaptation resulted in a smaller initial transient during second adaptation compared to a short abrupt protocol, both in the model and data. In this manuscript, we have commented mainly on qualitative results that are robust to model and protocol parameter choice and have not attempted quantitative fits. However, some other results — such as the relative ordering of savings between 'abrupt' and 'extended gradual' in this figure – are dependent on details of the model or experiment. One can change this ordering either by changing model parameters, or by changing the 'extended' duration of the 'extended gradual' protocol. Such sensitivity to experimental parameters is a falsifiable prediction that can be tested by further experiment. Source data are provided as a Source Data file.

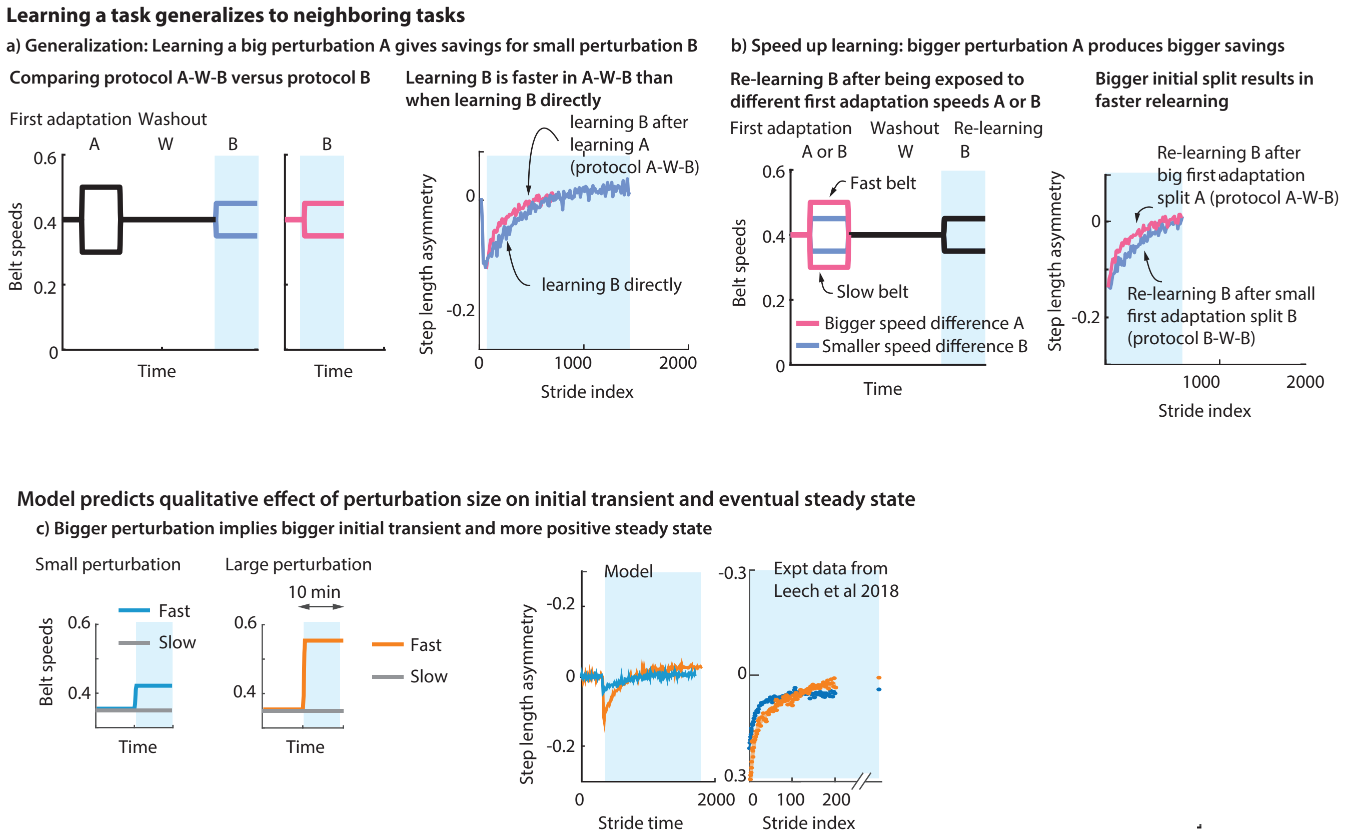


**Supplementary Figure 6. Generalization to neighboring tasks and size dependence of transients. a)** Learning a new task generalizes to neighboring tasks. Learning a larger split A and then learning a smaller split B with tied-belt washout W in between (protocol A-W-B) results in a faster learning transient for task B rather than learning B directly (protocol B), as seen in experiments[9]. Light blue shaded regions indicate adaptation regions for whom behaviors are compared. **b)** Learning a larger split A and then learning a smaller split B, with a tied-belt washout W in between (protocol A-W-B) can result in faster re-learning transient for B than learning B first and then re-learning B with a washout in between (protocol B-W-B), as in experiments[9]. These phenomena are due to the interpolative function approximation properties of the memory and constitute savings from experience with neighboring tasks. **c)** The model is able to capture, qualitatively, how the initial transient and the eventual steady state change based on the size of the belt-speed splits[9]. Source data are provided as a Source Data file.

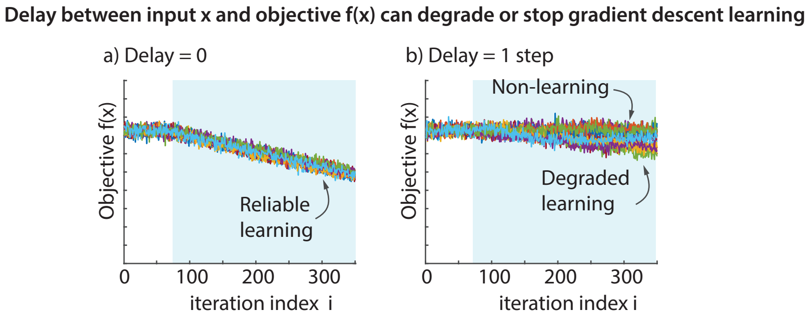


**Supplementary Figure 7. Delay degrades or stops adaptation.** Exploration-based gradient descent to optimize an objective function $f(x)$ with and without delay. Blue shaded region is when the optimization process is turned on. **a)** Without delay between $x$ and $f(x)$, the gradient is estimated reliably and gradient descent proceeds to reduce the function value. Twenty different optimizations are shown from the same initial condition, and same levels of exploratory and sensory noise levels. **b)** With a delay of one step between the input $x$ and function values $f(x)$, learning via gradient descent is completely stopped in some trials and is severely degraded in other trials. See *Supplementary Methods*. This delay-based degradation of learning is largely independent of the nominal energy gradients. Source data are provided as a Source Data file.

| Figure | Quantities compared | $t$(df) | $p$ | Cohen's $d$ | Paired | Tailed | Non-parametric $p$ | Normality |
|---|---|---|---|---|---|---|---|---|
| 5 | Model: time constants of first and second adaptation | 21.8(9) | $< 0.001$ | 9.62 (5.95, 16.18) | yes | yes | 0.002 | yes |
| 6a | Model: initial transient in A in TATATA vs TATBTA | 0.244(18) | 0.810 | 0.104 (-0.74, 0.94) | no | no | 0.678 | yes |
| 6a | Model: early change in A in TATATA vs TATBTA | -0.114(15) | 0.911 | -0.049 (-0.89, 0.79) | no | no | 0.910 | yes |
| 6b | Model: rate constant of B and A in TBA | -1.42(7) | 0.200 | 0.066 (-0.21, 0.048) | yes | no | 0.250 | yes |
| 6b | Experiment: rate constant of B and A in TBA | -0.363(8) | 0.726 | -0.11 (-0.70, 0.49) | yes | no | 1 | yes |
| 6b | Model: initial transient of B and A in TBA | -37.4(7) | $< 0.001$ | -18.1 (-32.9, -10.6) | yes | no | 0.008 | yes |
| 6b | Experiment: initial transient of B and A in TBA | -3.36(8) | 0.005 | -0.98 (-2.10, -0.25) | yes | no | 0.006 | yes |
| 7a | Model: after-effect noisy and no noise | 2.86(11) | 0.0154 | 1.13 (0.28, 1.96) | no | no | 0.001 | yes |
| 7a | Experiment: after-effect noisy and no noise | -2.41(14) | 0.0151 | -1.14 (-2.14,-0.11) | no | no | 0.0249 | yes |
| 7b | Model: gradual no-noise and gradual noisy | 3.59(12) | 0.0035 | 1.42 (0.53, 2.28) | no | no | 0.0051 | yes |
| 7b | Model: gradual no-noise and abrupt no noise | 11.9(22) | $< 0.001$ | 4.68 (3.09, 6.24) | no | no | $< 0.001$ | yes |
| 7b | Model: gradual noisy and abrupt no noise | 0.16(12) | 0.875 | .063 (-0.71, 0.84) | no | no | 0.371 | yes |

**Supplementary Table 1. Details of statistical comparisons.** Column 1 refers to the figure number in the main manuscript, in which the corresponding data are plotted. Details reported in columns 3, 4, 6, and 7 are for $t$-tests: $t$(df) indicates the $t$ statistic, with the number of degrees of freedom for the $t$ test in parenthesis; $p$ refers to the $p$ value for the $t$-test. Tailed $t$-tests are always in the direction of difference predicted by the model. Normality of samples was tested and we found that the null hypothesis of normality was not rejected at the 5% level for any of the samples. In addition to $t$-tests, $p$ value for non-parametric tests that do not rely on normality are provided. In every occurrence, the non-parametric test agreed with the t-test at the 0.05 significance level. See *Supplementary Methods* for more details.

### Supplementary Methods

#### Biped model, feedback controller, learning, and parameters

We now provide further mathematical details for the biped model[10], controllers, and learning.

**Equations of motion.** When the foot is fixed in an inertial frame, the inverted pendulum phase of inverted pendulum walking (Supplementary Figure 1a-b) has the following equations of motion:

$$m\ell^2\ddot{\theta} + mg\ell\sin\theta = \tau, \tag{1}$$

where $\theta$ is the stance leg angle from the vertical, $g$ is the acceleration due to gravity, $\ell$ is the leg length, $m$ is the point mass body, and $\tau$ is an ankle torque; clockwise and rightward are positive. These equations apply whether the inverted pendulum walking happens overground or on a treadmill, as long as the treadmill belt holds constant speed. A purely forward force $F$ on the center of mass, such as from an external device or the inertial forces from belt acceleration, is achieved by having $\tau = F\ell\cos\theta$. Thus, when the belt is accelerating forward with acceleration $a_{\text{belt}}$, the equation of motion is:

$$m\ell^2\ddot{\theta} + mg\ell\sin\theta = -m\,a_{\text{belt}}\,\ell\cos\theta. \tag{2}$$

**Push-off and heel-strike.** During the step to step transition (Supplementary Figure 1b-c), push-off happens first. The leg direction $\hat{n}$ is initially perpendicular to the body velocity $\bar{v}$ with respect to the current stance belt just before push-off. During push-off, if the push-off impulse is positive, the velocity is such that $\hat{n} \cdot \bar{v} \geq 0$, so that push-off only does positive work. If the push-off impulse magnitude is $I_{\text{push-off}}$, the post-push-off velocity in the push-off belt frame is $\bar{v} + \Delta\bar{v} = \bar{v} + I_{\text{push-off}}\,\hat{n}/m$. The positive work performed by the leg is equal to the kinetic energy change in the push-off belt frame, which is also equal to $m(\Delta v)^2/2$, because of pythagorean theorem[11]. This quantity is equal to the work performed by a telescoping leg.

The analysis of heel-strike is the reverse of push-off. After heel-strike, the body velocity is again perpendicular to the new stance leg, now on the heel-strike belt. If the body velocity just before heel-strike in the heel-strike belt frame is $\bar{v}$ and the heel-strike leg direction is now $\hat{n}$, the post-heel-strike velocity in the heel-strike belt frame is $\bar{v} + \Delta\bar{v} = \bar{v} - (\bar{v} \cdot \hat{n})\hat{n}$. Here, the second term $(\bar{v} \cdot \hat{n})\hat{n}$ is the component of pre-heel-strike velocity along the heel-strike leg, which is entirely lost upon heel-strike. The leg only performs negative work during such a heel-strike. The value of this negative work equals the change in kinetic energy across the heel-strike, computed in the frame of the belt onto which heel-strike happens, given by: $-m(\bar{v} \cdot \hat{n})^2/2$.

**Metabolic energy cost.** The metabolic energy cost $E_{\text{step}}$ for each step is a sum of two terms, a stance leg cost $E_{\text{stance}}$ and a leg swing cost $E_{\text{swing}}$. That is,

$$E_{\text{step}} = E_{\text{stance}} + E_{\text{swing}}. \tag{3}$$

The stance leg cost captures the metabolic cost of the mechanical effort during stance, and we set it equal to

$$E_{\text{stance}} = b_{\text{pos}} W_{\text{pos}} + b_{\text{neg}} |W_{\text{neg}}|, \tag{4}$$

where $W_{\text{pos}}$ and $W_{\text{neg}}$ are, respectively, the push-off positive work and the heel-strike negative work, and $b_{\text{pos}} = 4$ and $b_{\text{neg}} = 0.83$ are reciprocals of the positive and negative work efficiencies[1,12,13]. For the leg swing cost, we initially considered two versions of the model: (1) a work-based swing cost, based on the mechanical work needed to move a point foot by a given distance in a given amount of time, starting and ending at the respective belt speeds; this cost used the same weighting of positive and negative work as for the stance cost[10,11,13,14]; (2) an empirical swing cost due to Doke and Kuo[15] where the cost scales with the typical force rates required for the point-mass to traverse a given distance in a given time[11,13,15]. We confirmed that the qualitative step length asymmetry adaptation behavior for the split-belt adaptation was similar for the two swing costs, and then for the rest of the manuscript, used the latter force-rate-related cost given by the following equation:

$$E_{\text{swing}} = c_{\text{swing}} \left| \frac{\Delta v}{\Delta t} \right| \cdot \frac{1}{\Delta t}, \tag{5}$$

where $\Delta t$ is with the swing duration and $\Delta v$ is the change in the foot speed from stance (when the foot moves with the belt) to swing (when the foot moves with speed needed to cover the distance between the foot placements within the swing duration). The multiplier for the swing cost $c_{\text{swing}} = 0.9$ was chosen to approximate the speed-step-length relationship in normal walking (Supplementary Figure 2).

The metabolic rate over a stride (two steps) is the metabolic cost divided by the stride time. For purely visualization purposes, from this stride-wise metabolic rate $\dot{E}_{\text{stride}}$, we predict what would be measured via indirect calorimetry $\dot{E}_{\text{VO2}}$, obtained by filtering the stride-wise metabolic rate with a first order linear process with a time-constant of 42 seconds (as in[16,17]). That is,

$$\dot{E}_{\text{VO2}} = \lambda \cdot (\dot{E}_{\text{stride}} - \dot{E}_{\text{VO2}}) \tag{6}$$

where $\lambda$ is the reciprocal of the time constant. Such predicted metabolic equivalents of indirect calorimetry is denoted as measured by VO2 in Fig. 2 in the main manuscript.

**Feedback controller.** The inverted pendulum walker has only two control variables for each step (Supplementary Figure 1d): the push-off impulse $I$ and the step length $d$, so that the control variable $u = [I; d]$. These two control variables are modulated based on deviations in body state to keep the biped stable. The feedback controller is of the form: $u = u_{\text{nominal}} + K(s - s_{\text{nominal}})$ or equivalently $u = a + Ks$, with $a = u_{\text{nominal}} - Ks_{\text{nominal}}$, where $s$ is the relevant body state and $K$ is a matrix of feedback gains. The state $s$ is composed of four variables: the body velocity in the belt frame, body velocity in the lab frame, the body position in the lab frame, and the sum of past body positions in lab frame, all at midstance, that is when the body is over the foot ($\theta = 0$). This controller has the structure of a discrete *PID* control, with feedback on position (proportional term), velocity (derivative term), and sum of positions (integral term). The feedback on the body position and the sum of past body positions help with station-keeping on the treadmill, that is, not drift off the treadmill. We use the discrete sum instead of the integral because the control is discrete and once-per step, rather than continuous. The parameters $u_{\text{nominal}}$, $K$, and $s_{\text{nominal}}$ for the default controller are obtained from steady walking data, so as to approximate the step to step maps, as in[3–5]. This fit has one additional input from those in[3,4] and uses slightly different relative weighting for matching the step to step map – but the differences are unimportant for the qualitative results herein. The specific gain parameters are in Supplementary Table 2.

By default, we use the state at mid-stance to select the next push-off and step length, because in most simulations here, there is no additional perturbation between midstance and heelstrike[3–5]. Just for the case where there are impulsive exoskeletal input, instead of basing the control on the mid-stance state, we allow the target push-off and step length on the next step to be continuously updated through the stance phase until the leg angle corresponds to the proposed step length. This continuous updating of the proposed control is performed so as to account for any additional perturbations between mid-stance and heel-strike. We perform this continuous updating so as to exactly conserve the overall step-to-step dynamics of the mid-stance based control: specifically, we continuously update the control based on the current angular velocity by computing the corresponding previous mid-stance angular velocity and then using the mid-stance-based controller to decide the next push-off and step length.

**Leg swing dynamics.** In the aforementioned model description, the swing legs had no explicit dynamics coupled to the rest of the body, but were directly controlled by the feedback controller by specifying the next foot placement. This is a simple modeling choice with a rich tradition of explaining a variety of locomotor phenomena[11,14,18]. Despite the lack of explicit leg-swing dynamics, the model has a leg swing cost that is computed from the distance and duration between foot placements, as mentioned in the previous paragraph on metabolic cost.

Simulating the addition of asymmetric foot mass requires explicit leg swing dynamics and control, so the dynamics have an additional degree of freedom in this case. For this situation, we use a minimal dynamical model of leg swing in which the leg swing is initiated by a hip impulse and then coasts passively until heel-strike, as suggested by classic leg swing EMG data[18,19]. This impulse is an additional control variable and the initial leg swing state is an additional state variable. The controller is chosen, as earlier, such that the model has the closest return map to the empirical human step to step dynamics at steady state. In this case, the detailed feedback gain parameters are different from that in Table 2 but the overall controlled locomotor dynamics are approximately the same by construction, so the controller is considered the same. Because the belt speed is fixed here, it is not essential to use controller gains for speed relative to the belt frame and the lab frame. The swing leg dynamics are coupled to the center of mass only through the foot placement[20] and thus the nominal gait for this biped with leg swing is identical to the pure inverted pendulum walking gait considered earlier without leg swing dynamics.

**Walking with exoskeleton torques.** All exoskeleton simulations used the inverted pendulum walking model as described above. For all the exoskeleton calculations considered here, the treadmill belt speed is fixed, so separated controller gains for speed relative to the belt frame and the lab frame Table 2 was not needed, but just the sum of their gains assigned to the speed relative to the belt is sufficient.

We conceived the exoskeleton as providing a torque about the foot, which — for this simple model — can help simulate forward pulls, slopes, ankle torques, and hip torques. We performed three types of exoskeleton simulations. First, we simulated walking with time-periodic torques[21,22], which simply used the equation 1 with the relevant torques; the torques used were zero everywhere, except for brief periods when the torque rose and fell linearly, resulting in a torque impulse (Figure 2D in the main manuscript). In this case, it is useful to for the learner to include the relative perturbation timing from the exoskeleton as a sensed input state in addition to the biped state.

Second, we simulated walking with the torque impulse applied at specific leg angles, thereby making the exoskeleton assistance phase-periodic (Supplementary Fig. 4). This was achieved by starting the torque impulse at the desired leg angle via event detection in MATLAB.

Finally, we simulated the exoskeleton with assistance depending on step frequency[23–25]. In this case, the exoskeleton torque was constant over a step, and this constant was a function of the step frequency over the previous two steps. This simulation also involved a period where the walker's step frequency is constrained by a metronome, with the prescribed frequency changing providing the walker broad experience on the energy landscape. We simulated this metronome-constrained walking by pre-computing the walking gait at different frequencies and interpolating between them when a particular metronome frequency is commanded; see figure 4 of the main manuscript to see that this approach was satisfactory.

**Parameters.** The physical and metabolic parameters of the biped, as well as the parameters of the feedback controller were chosen independently of the adaptation experiments. Instead, they were fixed by using calculations from prior studies on the human walking steady state (Supplementary Figure 2)[3–5,18]. Aside from these parameters, the reinforcement learner has two parameters, the gradient descent learning rate $\alpha_g$ and the finite memory size $N_g$ used by the gradient estimator. The memory mechanism has two parameters: a learning rate for moving toward the memory $\alpha_m$ and a learning rate for updating the memory $\alpha_{\text{mf}}$ toward the current controller. Unless otherwise stated, we usually used the following values for these parameters: $\alpha_g = 1.2 \times 10^{-4}$, $N_g = 30$, $\alpha_m = 0.01$, and $\alpha_{\text{mf}} = 0.03$, all in non-dimensional terms. These values were not tuned based on adaptation data, but instead were selected to simply demonstrate that the model predicts the many qualitative phenomena illustrated in the main manuscript figures 2, 4, 5, 6, and 7and Supplementary Figure 9). We confirmed that each of these parameters can generally be changed by at least 20% in each direction (and in many situations by a bigger factor) while retaining the general qualitative features of the predictions. The exploratory noise, in the absence of sensory noise was chosen to be $2 \times 10^{-3}$ in the non-dimensional variables; the results are completely insensitive to this size in the linear regime of the dynamics, and the size primarily matters in comparison to the sensory noise (see Figure 4d in the main manuscript). The baseline sensory noise was multiplicative and only in the energy measurement: this baseline error standard deviation was $10^{-4}$ (or 0.01%) times mean energy). In Figure 4d of the main manuscript, except for this baseline case, the other noise condition cases had both multiplicative and additive noise. The additive noise standard deviation (in non-dimensional units) was taken to be the same value as the multiplicative noise scaling paramater. The noise range for was chosen to demonstrate how too much sensory noise may overwhelm the exploratory variability. For Figure 3of the main manuscript, we used additional sensory noise on the forward velocity, which is used by the feedback controller being tuned. Overall, because the experimental protocols differ widely even with a single paradigm such as split-belt walking and not all experimental details are typically published (e.g., accelerations when treadmill speeds are changed), we focus on predicting the qualitative features of the results rather than pursue a quantitative fit of the model to data or capturing the quantitative details of experimental protocols.

**Computational infrastructure.** The analysis was implemented in the widely-used scientific language MATLAB (version 2022a). Differential equations were simulated using `ode45`, with the event detection option to detect heel-strike and switch to the next step. For performing optimization calculations for fitting to data (not for simulating the 'learning'), we used `fmincon` in MATLAB. Simulation of learning is implemented as a stride-by-stride for-loop, in which the current control parameters are used to simulate one stride (two steps) and then the control parameters and the memory are changed according to the learning model outlined herein. Please code associated with this manuscript, LocAd[26], available without restrictions in a public repository.

#### Further experimental details

For experimental work, we used the Vicon motion capture system with Vicon Nexus 1.x, Bertec instrumented split-belt treadmill, controlled from a MATLAB interface to automatically change speeds. As part of this study, we performed two split-belt walking experiments, involving a total of three conditions, with a total of 25 participants. The two experiments had 16 and 9 participants, and our participant number per condition was shown to be sufficient in prior split-belt studies for null hypothesis testing of difference in means (Torres-Oviedo and Bastian 2012, Malone Vasudevan and Bastian 2011). The resulting p-values were either less than 0.02 when 'significant' with a threshold of 0.05, or were greater than 0.3, so clearly insignificant. There were no marginal cases. No data were excluded from the analyses.

Statistical comparisons were performed for results in Figures 5, 6, 7 of the main manuscript. Normality of samples were tested using the Kolmogorov-Smirnov test, with a null hypothesis of normality and significance threshold for rejecting normality at the 0.05 level. The default statistical tests are $t$-tests, paired or unpaired as indicated when results are reported, allowing for unequal variances. Non-parametric tests were performed to augment the parametric t-tests: for such non-parametric tests, when the test is paired, we use a Wilcoxon signrank test and when the test is unpaired, we use a Wilcoxon ranksum test, also termed the Mann-Whitney $U$ test. The non-parametric test always agreed with the parametric tests at the 0.05 threshold of significance. Model-based qualitative results reported in other figures are essentially deterministic and no statistical tests are necessary for these. The data were anonymized before analysis and the investigators were thus effectively blinded to group allocation (which was random). Because of the objective nature of the analyses, such blinding is not relevant to the interpretation of the results.

#### Modulating memory use to not degrade via gradient descent

Consider the performance metric or objective function $J(p)$ to be minimized and the following learning rule that combines gradient descent and progress toward memory:

$$p_{i+1} = p_i - \alpha_g \nabla_p J + \alpha_m (p_{\text{memory}} - p_i). \tag{7}$$

Here, $\nabla_p J$ is the gradient of the objective $J$ with respect to the variable $p$, when $p = p_i$. The stored motor memory $p_{\text{memory}}$ need not necessarily be the correct minimum of the performance metric. Here, we first show why $\alpha_m$ not being a constant is necessary, if this procedure must converge to an extremum of the objective. Next, we show that modulating $\alpha_m$ via a truncated cosine tuning is sufficient to allow convergence to an extremum of the objective.

**Necessary condition.** Assume $\alpha_m$ is constant. Fixed points $p^*$ of equation 7 are given by when $p_{i+1} = p_i = p^*$. So, fixed points $p^*$ satisfy: $p^* = p^* - \alpha_g \nabla_p J(p^*) + \alpha_m (p_{\text{memory}} - p^*)$, or

$$\alpha_g \nabla_p J(p^*) = \alpha_m (p_{\text{memory}} - p^*). \tag{8}$$

| Feedback gain parameter $K_{ij}$ | Value | Feedback gain parameter $K_{ij}$ | Value |
|---|---|---|---|
| $K_{11}$ | -0.5211 | $K_{21}$ | 0.2793 |
| $K_{12}$ | -0.0859 | $K_{22}$ | -0.0290 |
| $K_{13}$ | -0.0073 | $K_{23}$ | 0.0020 |
| $K_{14}$ | -1.0559 | $K_{24}$ | -0.0750 |

**Supplementary Table 2.** Parameters of the default feedback controller. Here, $K_{jk}$ is defined as $\Delta u_j/\Delta s_k$, where $u$ is the control and $s$ is the state. For the control variable, $u_1$ is the push-off and $u_2$ is the swing leg angle at touch down. For the state variable $s_1 = \dot{y}_{\text{body/belt}}$, $s_2 = y_{\text{body/ground}}$, $s_3 = \sum y_{\text{body/ground}}$, $s_4 = \dot{y}_{\text{body/ground}} - \dot{y}_{\text{body/belt}}$. All numbers are non-dimensional.

Thus, the only situation this procedure converges to an extremum of the objective at which the gradient equals zero ($\nabla_p J(p^*) = 0$) is in the special case where $p_{\text{memory}}$ equals the optimum. For generic $p_{\text{memory}}$ not equal to the optimum, the system will converge to a $p^*$ such that the step along the negative gradient exactly cancels the step toward the memory, as in equation 8. Thus, $\alpha_m$ cannot be constant and its modulation in *some* manner is necessary. This necessary modulation is not unique, and in the following paragraph, we prove the sufficiency of one such modulation.

**Sufficient condition.** Assume that $\alpha_m = \alpha_0\, h(\theta_{gm})$, where $\alpha_0$ is a constant and $\theta_{gm}$ is the angle between the negative gradient $-\nabla_p J$ and the current direction toward memory: $p_{\text{memory}} - p$. Say, the function $h(\theta_{gm}) = 0$ when $|\theta_{gm}| > 90$ degrees and $h(\theta_{gm}) \neq 0$ when $|\theta_{gm}| < 90$ degrees. One example of such a function $h$ is a truncated cosine tuning, with $h(\theta_{gm}) = 0$ when $|\theta_{gm}| > 90$ degrees.

First, given the assumptions on $h(\theta_{gm})$, the iterations always proceeds along a descent direction, that is one that reduces the objective. This is true because:

1. when $|\theta_{gm}| \geq 90$, we have $\alpha_m = 0$, so the algorithm (equation 7) reduces to gradient descent and thus always steps along a descent direction;

2. when $|\theta_{gm}| < 90$, the step toward memory $(p_{\text{memory}} - p^*)$ has a positive component along the negative gradient by assumption of $\theta_{gm}$. Thus, $(p_{\text{memory}} - p^*)$ is a descent direction, and so the combined gradient descent step and the step toward memory in equation 7 is always a descent direction.

Second, given the assumptions on $h(\theta_{gm})$, we now show that $p_{\text{opt}}$ with $\nabla J = 0$ is a fixed point of the algorithm. Say there exists an optimal point $p_{\text{opt}}$ where $\nabla J = 0$ and $p_{\text{opt}} \neq p_{\text{memory}}$. Then, there exists an $\varepsilon$ neighborhood $S$ around $p_{\text{opt}}$, not containing $p_{\text{memory}}$ which can be divided into two disjoint sets $S_1$ and $S_2$, with $p_{\text{opt}}$ being at the boundary separating $S_1$ and $S_2$a – with the two sets defined as follows. The set $S_1$ is defined so that any point $p \in S_1$ satisfies $|\theta_{gm}| \geq 90$ degrees, so that when the current iterate falls in this region, the learning algorithm defaults to regular gradient descent because $\alpha_m = 0$. Thus, when the iterates $p_i$ remain in $S_1$, $p^* = p_{\text{opt}}$ is an accumulation point of the iteration. The complementary set $S_2$ is defined such that any point $p \in S_2$ satisfies $|\theta_{gm}| < 90$ degrees, but this set cannot contain any fixed points because $|\theta_{gm}| < 90$ is incompatible with the fixed point equation $-\alpha_g \nabla_p J(p) + \alpha_m(p_{\text{memory}} - p_i) = 0$. This can be seen by taking the dot product of this fixed point equation with $-\nabla_p J(p)$ and noting that the resulting scalar left hand side is non-negative and the right hand side is negative by definition. Thus, the only fixed point of the algorithm in the neighborhood of $p_{\text{opt}}$ is $p^* = p_{\text{opt}}$, in the set $S_1$.

Thus, modulating the $\alpha_m$ using a truncated cosine tuning is sufficient to ensure that (1) the algorithm always reduces the objective and (2) a fixed point of the descent algorithm is the optimal point $p_{\text{opt}}$. Of course, what we have shown does not guarantee convergence to a local minimum, which we leave for an examination in a future study.

#### Numerical experiments to show that delayed device response can degrade or stop optimization

In a number of exoskeleton adaptation experiments[23,24,27,28], the assistance or resistance provided by the exoskeleton on one step depends on what the human did on the previous step (e.g., the previous step period or step width). Some treadmill adaptation studies also had a similar protocol[29], the treadmill speed is based on previous step periods. At least in some of these studies[23,24,27,28], many subjects did not initiate adaptation despite the obvious energetic advantage to such adaptation. Here, we argue that this non-adaptation could be due to the temporal delay between human action and the energetic reward or punishment. If the human nervous system is performing energy optimization based on correlating motor actions on a particular step and their energetic consequences, we may expect such an energy optimization to fail when action and reward are separated. Indeed, there is some precedent for delay affecting motor control. However, the story is a bit more subtle and whether or not adaptation happens depends on the exploratory optimization algorithm and the exploration. Let us consider two example algorithms, including the algorithm we used. Say the goal is to find $x$ to minimize the function $f(x)$ and with $x_0$ being the initial guess for the optimum. For simplicity, let us assume that the system has no additional dynamics.

**Algorithm 1. Simplest hill climbing.** While the term 'hill climbing' has sometimes been associated with gradient descent (more precisely, gradient ascent), here we refer to an even simpler local search algorithm[23]. Say the best guess for the optimum at step $i$ is $x_i$ with function value $f(x_i)$. The optimizer then evaluates the function at $\hat{x}_{i+1} = x_i + \mu_i$, that is, evaluates $f(x_i + \mu_i)$, where $\mu_i$ is exploratory noise having Gaussian distribution and with the usual assumptions of Gaussian and independent from the previous $\mu$'s. The next best guess $x_{i+1}$ of the optimum is decided by the following rule:

$$x_{i+1} = x_i + \mu_i \text{ if } f(x_i + \mu_i) < f(x_i) \tag{9}$$

$$x_{i+1} = x_i \text{ if } f(x_i + \mu_i) \geq f(x_i). \tag{10}$$

That is, make a new guess; if the new guess improves the objective, go there; if not, stay where you are. It is well known that this algorithm goes toward local minima under normal circumstances. But if there is a 'delay' unbeknownst to the algorithm, for instance, wherein $f(\hat{x}_{i+1})$ is actually $f(\hat{x}_i)$, then decisions to accept will be based on incorrect function values and it is clear that this algorithm will not be optimizing.

**Algorithm 2. Gradient descent with isotropic exploration** This is the gradient descent that we have used in this study, in which the gradient is estimated via linear regression between input and output over the past $N_{\text{gradient}}$ steps. Here, the exploratory noise is isotropic, so all directions are equally likely. Without any delay between action $x$ and function evaluation $f$, the algorithm reliably goes toward a local minimum. When there is a delay, the algorithm may sometimes go toward the optimum, thus 'adapting' and sometimes not make any progress, thus 'not adapting', even for the exact same values of all parameters, as shown in Fig. 10of the main manuscript. The reason the algorithm can sometimes go toward the optimum despite a delay is because the gradients on successive steps are correlated or persistent. Because successive steps are correlated, despite the function value mismatch, the algorithm can infer a reasonable descent direction as the gradient estimate, even if the gradient estimate may itself be biased.

### References


1. Srinivasan, M. & Ruina, A. Computer optimization of a minimal biped model discovers walking and running. *Nature* **439**, 72–75 (2006).
2. Srinivasan, M. Fifteen observations on the structure of energy-minimizing gaits in many simple biped models. *J. R. Soc. Interface* **8**, 74–98 (2011).
3. Wang, Y. & Srinivasan, M. Stepping in the direction of the fall: the next foot placement can be predicted from current upper body state in steady-state walking. *Biol. Lett.* **10**, 20140405 (2014).
4. Joshi, V. & Srinivasan, M. A controller for walking derived from how humans recover from perturbations. *J. Roy. Soc. Interface* **16**, 20190027 (2019).
5. Seethapathi, N. & Srinivasan, M. Step-to-step variations in human running reveal how humans run without falling. *ELife* **8**, e38371 (2019).
6. Bertram, J. & Ruina, A. Multiple walking speed-frequency relations are predicted by constrained optimization. *J. theor. Biol.* **209**, 445–453 (2001).
7. Minetti, A. & Alexander, R. A theory of metabolic costs for bipedal gaits. *J. Theor. Biol.* **186**, 467–476 (1997).
8. Roemmich, R. T. & Bastian, A. J. Two ways to save a newly learned motor pattern. *J. Neurophys.* **113**, 3519–3530 (2015).
9. Leech, K. A., Roemmich, R. T. & Bastian, A. J. Creating flexible motor memories in human walking. *Sci. Rep.* **8**, 1–10 (2018).
10. Srinivasan, M. Fifteen observations on the structure of energy-minimizing gaits in many simple biped models. *J. R. Soc. Interface* **8**, 74–98 (2011).
11. Seethapathi, N. & Srinivasan, M. The metabolic cost of changing walking speeds is significant, implies lower optimal speeds for shorter distances, and increases daily energy estimates. *Biol. Lett.* **11**, 20150486 (2015).
12. Ruina, A., Bertram, J. E. & Srinivasan, M. A collisional model of the energetic cost of support work qualitatively explains leg sequencing in walking and galloping, pseudo-elastic leg behavior in running and the walk-to-run transition. *J. theoretical biology* **237**, 170–192 (2005).
13. Seethapathi, N. *Transients, Variability, Stability and Energy in Human Locomotion*. Ph.D. thesis, The Ohio State University (2018).
14. Handford, M. L. & Srinivasan, M. Sideways walking: preferred is slow, slow is optimal, and optimal is expensive. *Biol. letters* **10**, 20131006 (2014).

**15.** Doke, J., Donelan, J. & Kuo, A. Mechanics and energetics of swinging the human leg. *The J. Exp. Biol.* **208**, 439–445 (2005).

**16.** Selinger, J. C. & Donelan, J. M. Estimating instantaneous energetic cost during non-steady-state gait. *J. Appl. Physiol.* **117**, 1406–1415 (2014).

**17.** Miyamoto, Y. *et al.* Dynamics of cardiac, respiratory, and metabolic function in men in response to step work load. *J. Appl. Physiol.* **52**, 1198–1208 (1982).

**18.** Kuo, A. A simple model of bipedal walking predicts the preferred speed–step length relationship. *J. Biomech. Eng.* **123**, 264–269 (2001).

**19.** Basmajian, J. V. Muscles alive. their functions revealed by electromyography. *Acad. Medicine* **37**, 802 (1962).

**20.** Garcia, M., Chatterjee, A., Ruina, A. & Coleman, M. The simplest walking model: stability, complexity, and scaling. *ASME J. Biomech. Eng.* **120**, 281–288 (1998).

**21.** Ahn, J. & Hogan, N. A simple state-determined model reproduces entrainment and phase-locking of human walking. *PloS one* **7**, e47963 (2012).

**22.** Ochoa, J., Sternad, D. & Hogan, N. Treadmill vs. overground walking: different response to physical interaction. *J. Neurophysiol.* **118**, 2089–2102 (2017).

**23.** Selinger, J. C., Wong, J. D., Simha, S. N. & Donelan, J. M. How humans initiate energy optimization and converge on their optimal gaits. *J. Exp. Biol.* **222**, jeb198234 (2019).

**24.** Selinger, J. C., O Connor, S. M., Wong, J. D. & Donelan, J. M. Humans can continuously optimize energetic cost during walking. *Curr. Biol.* **25**, 2452–2456 (2015).

**25.** Simha, S. N., Wong, J. D., Selinger, J. C., Abram, S. J. & Donelan, J. M. Increasing the gradient of energetic cost does not initiate adaptation in human walking. *bioRxiv* (2020).

**26.** Seethapathi, N., Clark, B. & Srinivasan, M. Locad: Code for 'exploration-based learning of a stabilizing controller predicts locomotor adaptation'. https://github.com/SeethapathiLab/LocAd (2024).

**27.** Simha, S. N., Wong, J. D., Selinger, J. C., Abram, S. J. & Donelan, J. M. Increasing the gradient of energetic cost does not initiate adaptation in human walking. *J. Neurophysiol.* **126**, 440–450 (2021).

**28.** Abram, S. J., Selinger, J. C. & Donelan, J. M. Energy optimization is a major objective in the real-time control of step width in human walking. *J. biomechanics* **91**, 85–91 (2019).

**29.** Snaterse, M., Ton, R., Kuo, A. D. & Donelan, J. M. Distinct fast and slow processes contribute to the selection of preferred step frequency during human walking. *J. Appl. Physiol.* **110**, 1682–1690 (2011).
